# Membrane bending energy selects for compact growth of protein assemblies

**DOI:** 10.1101/2025.08.08.669413

**Authors:** Yue Moon Ying, Margaret E. Johnson

## Abstract

Remodeling of cell membranes into vesicles is essential for receptor transport into cells and viral escape from infected cells. Membranes must be forced into these highly curved vesicles, and this is primarily driven through a structured assembly of multiple, multivalent interacting protein subunits forming a lattice. Lattice assembly from these subunits is a stochastic process, and intermediate structures formed during growth can vary in both structure and stability. Here we show that the membrane bending energy cost per protein rises significantly when remodeling is driven by lattice intermediates that deviate from compact, ideal spherical structures. We use a continuum membrane mechanics model coupled to lattice intermediates assembled from stochastic rigid-body simulations of HIV-1 Gag lattice assembly to quantify the bending energy as it systematically varies with lattice eccentricity. Our results show that highly eccentric lattices induce a higher bending energy cost because the lattices still deform the membrane into an approximate spherical cap, but the radius of the cap is larger due to the imperfect lattice geometry. These quantitative trends are also nearly independent of the density of links to the membrane, emphasizing the importance of the lattice perimeter shape instead.

Rescaling thus recovers an approximately universal bending energy cost when evaluated relative to the circumscribing sphere of the lattice intermediates. These results show that assembly pathways coupled to membrane remodeling face much stronger selection pressure for highly compact growth compared to solution assembly pathways due to bending energy costs and provide a tool to characterize these pathways during processes like viral budding and endocytosis.

## 1. Introduction

Membrane remodeling driven by multi-protein assemblies is essential for diverse cellular processes such as viral budding, endocytosis, and cell division[REF]. Reshaping the two-leaflet membrane bilayer requires work to overcome the preferred flat equilibrium of a bilayer with a symmetric leaflet composition. Individual proteins can induce changes from this flat curvature preference by inserting amphipathic helices into a single leaflet. Asymmetry in lipid composition between leaflets can further bias the preferred membrane curvature, as occurs during budding of the HIV virion^1,2^, with local reduction in membrane surface tension ^3^. However, these contributions are not sufficient for large-scale membrane reshaping or formation of spherical membrane vesicles in cells, which rely on the self-assembly of multi-valent protein structures to impart relatively rigid reshaping forces. The budding of the HIV-1 virion out of the cell plasma membrane requires the assembly of the HIV Gag polyprotein (group-specific antigen ^4^) into an incomplete spherical lattice with a tri-hexagonal structure containing ∼2500 Gag monomers ^5-10^. The self-assembly of the Gag lattice and the remodeling of the membrane are coupled, as the Gag is observed to bind to the plasma membrane as monomers or small oligomers, not a complete lattice^11,12^. Although several studies have used computation to interrogate how multi-protein assemblies induce membrane remodeling via clathrin cages^13^, colloids^14^, and filaments^15^, our focus here is on the inverse cause-and-effect: how will the mechanical cost of membrane bending impact self-assembly pathways via energetic selection? Using a continuum membrane model linked to stochastically assembled rigid Gag lattices from reaction-diffusion simulations^16^, we here compare how deformations induced by highly regular and compact lattices versus irregular or eccentric lattices containing the same monomer numbers produce a much lower bending energy cost. The importance of the radially ‘ideal’ lattice structures on the bending energy cost has not been systematically tested before, and our result predicts a strong energy penalty on any self-assembly pathway that deviates significantly from ideal, radial growth.

With computer simulations of coarse-grained molecular subunits, one can track all intermediates along assembly pathways and compare their structure and energetics^16-21^, exceeding the spatial or temporal resolution of imaging or in vitro experiments^22,23^. The structures of multi-protein assemblies is critical in controlling the shape and thus bending energy of the membrane, driving vesicles with curvature matched to the assembly radius^13,24^, tubules^25^, or neck-like constrictions^26^. Given that most of these protein assemblies (including the Gag lattice^22,27,28^) can also form in solution, decoupled from membrane bending, we ask here how the energetic cost of membrane bending might substantially narrow assembly pathways, culling out structures accessible in 3D^16^. Because the assembly of the Gag lattice on the membrane spans several hundred nanometers and 5∼25 minutes ^29,30^, with >10^3^ Gag monomers and >10^5^ lipids, any start-to-finish simulations of the coupled self-assembly with *dynamic* membrane remodeling are inaccessible to current tools^31^. Here we therefore decouple the dynamics of reaction-diffusion simulations of self-assembly^16^ from the membrane remodeling dynamics and use energy minimization of the membrane. Alternatively, ultra coarse-grained molecular dynamics (CG-MD) simulations have successfully assessed other sub-steps of coupled assembly and membrane budding pathways. For the HIV Gag system, CG-MD simulations revealed how inducing local membrane curvature enhanced Gag lattice assembly and growth ^19^. Membranes coated with CG-MD models of BAR-domain proteins induce dramatic tubulation of liposomes^25^, and simulations of a viral dodecahedron revealed lipid microdomains were needed to drive fully complete assembly and budding^24^. Pre-assembled filaments on membranes can drive membrane tubulation and scission during cell division ^32,33^ using a coarse-grained one-particle-thick membrane model ^34^, and simulations with the CG-MD Cooke model^35^ showed that clustered proteins can drive membrane remodeling via curvature-induced interactions^36^. The primary limitation of CG-MD models is their expense, where extensions to the full budding of the HIV-1 Gag lattice will require systems with membranes several-fold larger than current studies, potentially requiring hybrid particle-based methods^37^. The membrane material properties such as bending modulus and tension are also more challenging to calibrate to experiment. Here we instead use a thin-film continuum model for the membrane^38,39^, which readily scales to micron lengths and is parameterized by material properties directly.

By abstracting the lipid membrane to an elastic curved surface, thin-film or single-layer continuum models provide substantial speed-ups compared to CG-MD (particle-based) membrane models, supporting much larger membrane simulations. With an energy function typically defined via the Helfrich-Canham-Evans Hamiltonian to describe the bending energy and tension energy of lipid bilayers ^40-42^, these models are more readily parameterized to changing material properties, albeit at the cost of capturing lipid heterogeneity and dynamics. Numerical implementations of continuum surface models use the finite element method (FEM) with various discretization methods ^43-45^; in contrast to discrete differential geometric methods^46 47^, here we use the subdivision limit-surface technique^48,49^ which allows for continuous and differentiable measurements of membrane curvature along the surface ^38^ (Fig 1). We previously showed that this model can recapitulate curvature sensing by amphipathic helices inserted to the bilayer with both one^39^ and two leaflets^50^, with excellent agreement to multiple experimental studies^50-52^. Continuum models coupled to densities of proteins are more efficient than particle-based protein models, revealing dramatic shape changes driven by dense protein fields^47,53-55^. However, structural variations induced locally by stochastically assembled structures are not observable. With particle-based protein models, deformations from cytoskeletal filaments^56,57^, lattice structures^13,58-60^, and flexible monomers^26,61^ predict how localized deformations couple to protein shape. Unlike these previous studies, here we set out to specifically quantify whether structural variations in assembled structures, physically accessible through stochastic 3D assembly^16^, represent realistic assembly pathways when coupled to a mechanical membrane. Furthermore, we can compare with theoretical estimates of membrane bending driven by spherical caps^14^, based on the size of the lattice-driven deformations in our numerical simulations.

**Figure 1.**
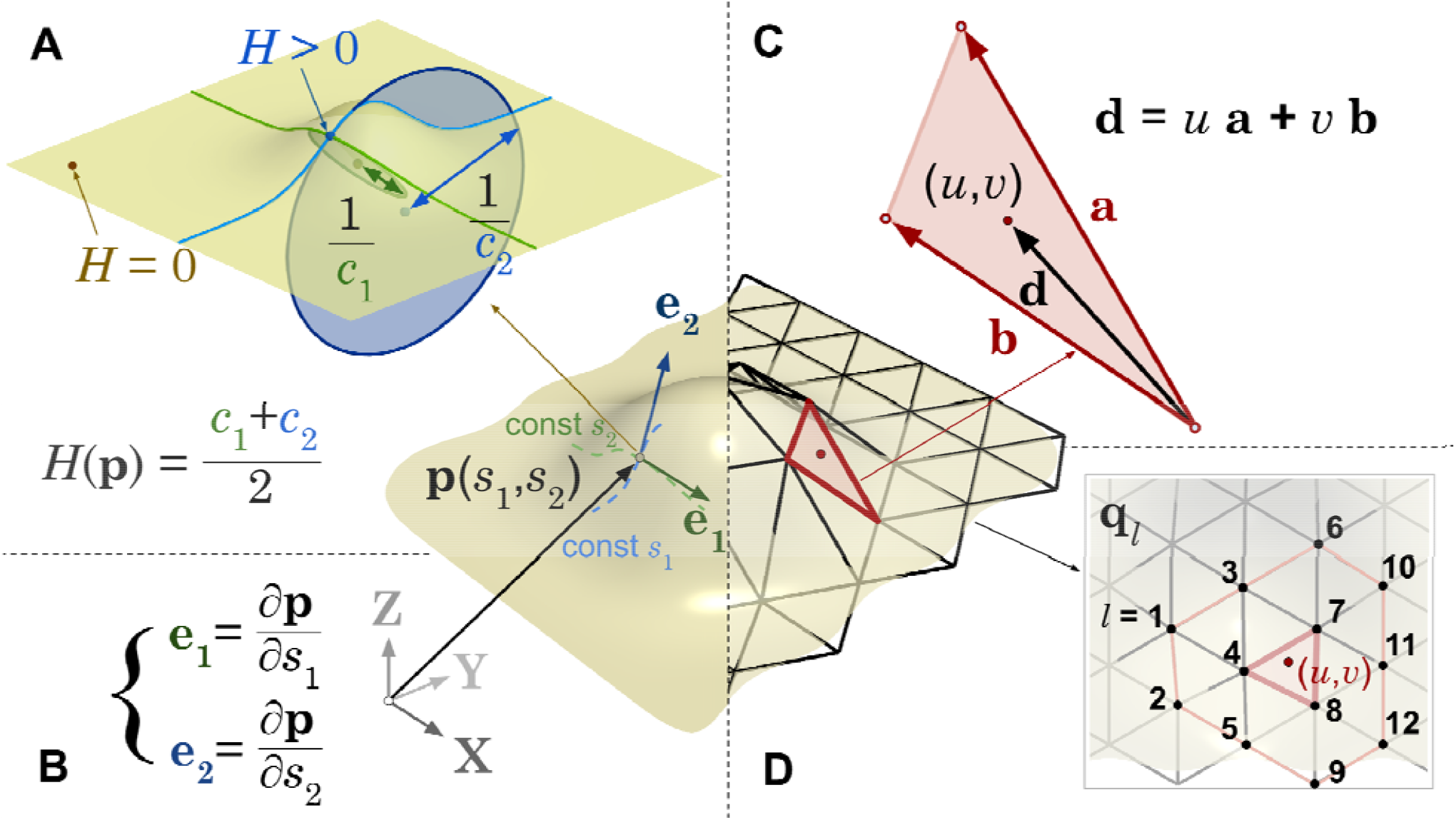
The continuum membrane model quantifies the membrane energy as it depends on shape due to variations in local curvature and total area, numerically represented using finite element methods. A) Mean curvature at some point on the limit surface is defined as the average of curvatures along two orthogonal tangent basis vector at this point (see Fig S1); B) Definition of tangent basis vectors **e**_1_,**e**_2_ at point on the limit surface via curvilinear coordinates (*s*_1_, *s*_2_); C) barycentric coordinate (*u, v*) defined inside an element triangle via linear combination of the edge vectors; D) Any point on the continual limit surface can be calculated with the coordinator vectors of the 12 neighboring vertices **q**_*l*_ (l enumerates the neighboring vertices) and barycentric coordinate (*u, v*) of the point on the triangle.

The HIV Gag immature lattice is an excellent system for studying the role of membrane mechanics on self-assembly pathways. It can be driven to assemble in solution given co-factors into structures that closely resemble the membrane-bound lattices^22,28^. From stochastic reaction-diffusion self-assembly simulations^62^ mimicking in vitro conditions, the Gag monomers initialized in the bulk (out-of-equilibrium) assemble into structures consistent with cryoET. However, the formation of complete structures^16^ and their remodeling dynamics^63^ is quite sensitive to the assembly conditions, producing a wide array of intermediate structures along assembly pathways with irregular or eccentric perimeters. Assembly co-factors are known to accelerate the kinetics of assembly in solution to improve yield^21^ and to avoid kinetically trapped intermediates that inhibit lattice completion in solution^16^. Because kinetically trapped states can persist for hours^64^, we reasoned that membrane localization and bending could provide another selection mechanism to ensure regular and robust growth to complete viral assemblies.

In this study, we first use rigid Gag lattices stochastically assembled in solution to quantify the bending energy cost associated with structures of varying regularity, despite having comparable numbers of Gag monomers. Comparison to theory indicates a substantial energetic penalty for irregular structures. This indicates the coupling of lattice assembly and membrane remodeling simultaneously would place energetic selection pressure on some structures over others, compared to in 3D. To more systematically confirm this observation and explain with theory, we generated highly-controlled lattices with controllable eccentricity (deviation from sphere to ellipse) and with varying density of membrane attachment sites. We show that the membrane bending energy is largely controlled by the overall shape and eccentricity of the lattice perimeter rather than by Gag attachment density. Finally, we re-scale our plots of membrane bending energy vs Gag monomer numbers to control for variations of eccentricity via the radius of the circumscribing cylinder around the maximal lattice dimension, producing a reasonably universal scaling of the bending energy as a function of this maximal radial lengthscale.

## II. METHODS AND MODELS

Our model systems combine a continuum surface model of a membrane with a rigid protein lattice, coupled together via harmonic bond potentials. We will assume here that because the protein lattice is rigid, its energy is fixed and unchanging. The membrane will reshape to minimize the energy imposed by the harmonic constraints coupling it to the protein lattice, and its own mechanics. Our continuum surface model has mechanics controlled by the Helfrich-Canham-Evans, or HCE ^40,42,65^ Hamiltonian. To evaluate the shape and energy of our continuum membrane as it adopts arbitrary topologies, we numerically represent it in 3D space using a Finite Element Method (FEM) that applies the limit surface method to smoothly parameterize the surface ^38^.

### II.Theory on the membrane and membrane-protein energetics

#### II.A.1 Background on the Energy Function describing the isolated membrane mechanics

Our continuum membrane model uses the HCE Hamiltonian *E*_tot_ (**p**,*A*) consisting of a membrane bending energy *E*_*B*_ and area constraint energy *E*_*A*_ (Eq. 1), which we re-define here for background. The membrane bending energy *E*_*B*_ quantifies the cost of bending dependent on a material parameter κ, the bending modulus, and the preferred or spontaneous curvature along the surface *C*_0_. It is defined as an integration over the whole membrane surface *S* of a quadratic penalty of local mean curvature *H*(**p**) measured at each point **p** on the curvilinear surface as it deviates from *C*_0_ (**p**) (Figure 1A). Here we set the spontaneous curvature to 0 nm^-1^ or flat (⩝**p** ∈*S,C*_0_(**p**)=0), which is true for bilayer membranes with symmetric leaflets. The local mean curvature 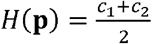, where C_1,2_ measure the principal surface curvatures, is therefore zero for a flat membrane. The bending modulus κ accounts for the free energy cost as the membrane deformation increases and is set to 83.4 pN⋅nm consistent with experimental measurements ^66,67^. Because κ is an emergent material property of the lipid membrane, unless in a system with phase separation ^68-70^, κ is constant throughout the membrane surface. The area constraint energy *E*_*A*_ is defined in terms of total membrane area *A* and equilibrium membrane area *A*_0_, which is the total membrane area if under zero tension. It is parameterized by the constant *μ*_*A*_ set to 250.0 pN/nm. In total we therefore have:

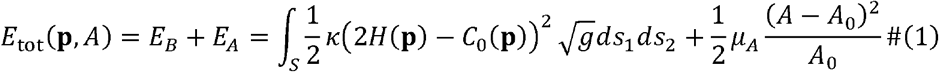

Where 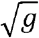 is the curvilinear factor required for area integration on a curved surface defined as the magnitude of the cross product of two tangent basis vector 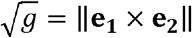 (see Fig 1B). When the spontaneous curvature is set to zero for the symmetric membrane, the bending energy simplifies to:

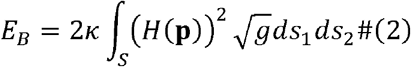

Tension τin this study is reported as the partial derivative of area constraint energy with respect to global area. Therefore, we can calculate tension based on area constraint energy from our simulations according to the following formula:

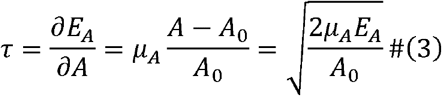

All parameters are defined along with the values used in Table S1, and for completeness, we also define all other variables of the theory/model in Table S2.

#### II.A.2. Energy function coupling the rigid lattice to the continuum membrane

The rigid body interaction model is a simplified model of the interaction between the protein lattice and the lipid membrane. The bond between the lattice and the membrane is simplified to a harmonic bond connecting the center of mass of a protein monomer (here the Gag protein) and the corresponding point on the limit surface of the control mesh vertex (Equation 4), which is initially set to the closest mesh vertex to the protein’s center of mass and periodically updated every fixed number of steps. The harmonic bond potential and force used in rigid body interaction model is defined as:

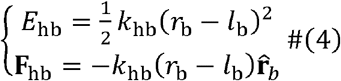

Where *E*_hb_ and *F*_hb_ are the energy and force exerted on the membrane due to the harmonic bond with parameters *k* _hb_ (the force constant) and *l*_b_ (the equilibrium bond length). *r*_b_ is the distance between the protein and the membrane, and 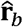 is the unit vector of the displacement vector pointing from scaffolding protein to the lipid. Although the actual interaction between HIV Gag protein and lipid membrane involves a series of hydrophobic and electrostatic interactions between Gag’s MA domain and PI(4,5)P_2_ ^73,74^, the bond length here describes a theoretical equilibrium distance between the center of mass of Gag protein and the membrane midplane. In the simulation, *l*_b_ is set to 9 nm.

We assume that the membrane is much softer than the protein lattice, which is a reasonable approximation for this study given estimated bending moduli of ∼10 fold higher for viral lattices ^75-77^. We also assume the links connecting the lattice to the membrane are effectively rigid as well to isolate all of the remodeling to the membrane surface. We therefore set *k*_hb_ to the very large value of 10^3^ pN/nm, to keep these bonds effectively rigid. In future work, both the protein lattice and the links would also be allowed to deform to minimize the system energy, as a more realistic representation of a protein-membrane system. This introduces additional parameters that are not necessary here, hence our simplifying assumptions.

The total membrane energy of the coupled membrane-lattice system will be the sum of the continuum membrane terms along with the contributions from the harmonic bonds connecting the surface to the lattice

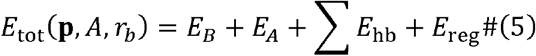

where here we note the addition of the purely numerical (not physical) regularization energy *E* _reg_,which serves to protect the triangular mesh from deforming during gradient descent to relatively uniform sizes and shapes of triangles for stable numerical integration ^38^ (Fig.1). *E*_reg_ should converge to near zero as the numerical minimization of the system energy maintain reaches its minimum and the forces also converge to zero. For analytical calculations, we therefore set *E*_*reg*_ = 0.

#### II.A.4. Analytical solutions to minimum energy structure of a membrane adhered to a spherical cap

For comparison with our numerical simulation results, we establish a series of theoretical approximations to the membrane bending energy induced by a perfectly spherical cap with a spherical radius of *r*_*M*_. Previous work has analytically derived the membrane shape function that aminimizes the energy of the lipid membrane upon attaching to a sphere, with and without membrane tension ^14^. Specifically, the attachment region is assumed be the same as the spherical surface. We follow the same approach as previous work by Deserno ^14^ but make one change, as we do not assume an infinite boundary condition; instead, we assume a finite boundary condition where the relaxation region becomes flat 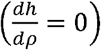 at a fixed distance *ρ*= *ρ*_*R*_ and the rest of the membrane with *ρ* > *ρ* _*R*_ is completely flat, as a more accurate representation of the finite simulated system size (see Fig. 2). The simplest cap model assumes that only the attachment region matters and the energy of the rest of the membrane is negligible. The bending energy of the attachment cap region *E*_*B*,cap_ is then:

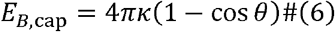

**Figure 2.**
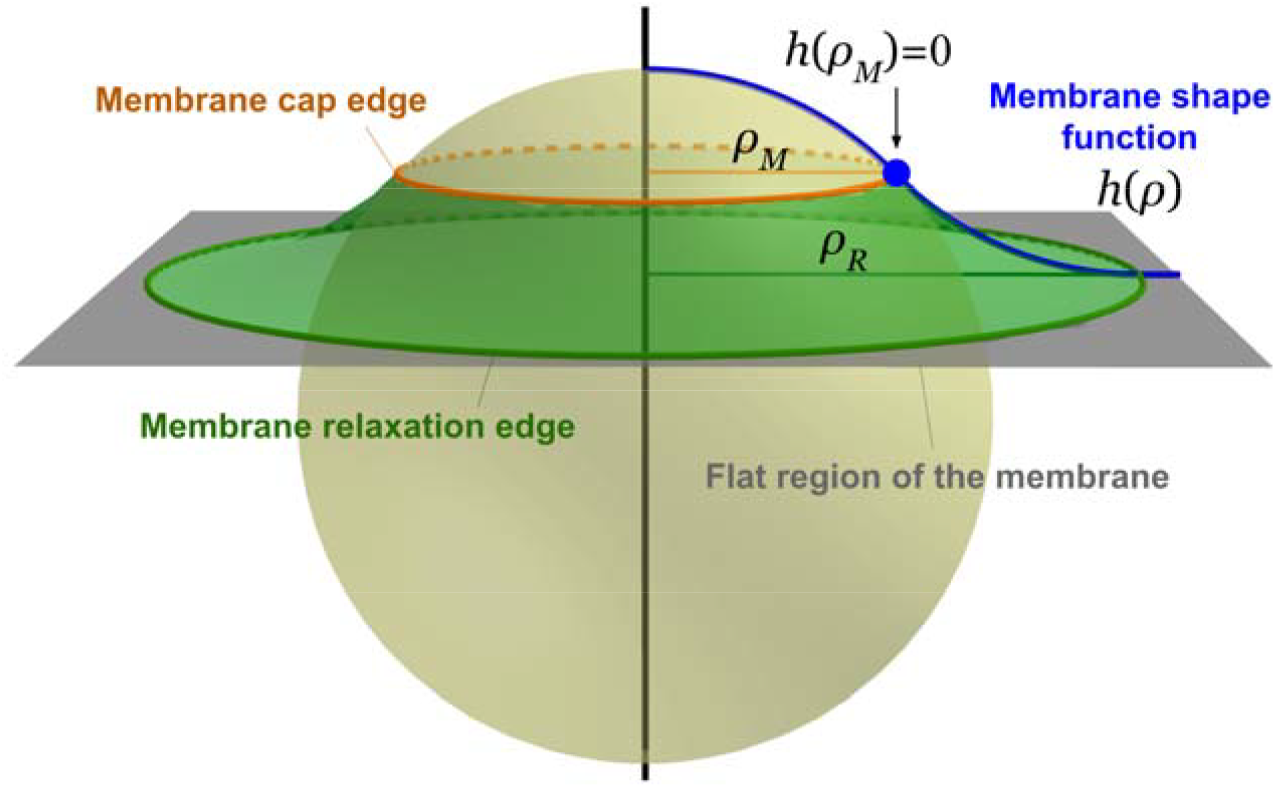
Theory can predict the bending energy of a membrane deformed by a sphere, including a continuous curvature relaxation. We illustrate here how the shape function that measures the membrane height at a radius of from the z-axis (black vertical line) is defined assuming a perfectly spherical indentation with azimuthally symmetric relaxation. The scaffolded spherical cap is delimited by the orange line at a radius of ρ_*M*_. The membrane at large distances is assumed to be flat (gray surface). The relaxation region in green captures the continuous change in curvature from the cap to the flat membrane at the radius ρ_*R*_. The membrane shape function captures these changes in curvature, as the theory assumes radial symmetry.

Where θ is the polar angle from the z-axis to the radii pointing to the end of the spherical cap.

Therefore, 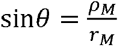 and *ρ*_*M*_ is the radial distances from the z-axis to the start of the relaxation function (see Fig 2, Fig S2). With no relaxation region, *ρ* _*M*_ is the radius where the sphere intercepts the flat plane. However, if we consider a patch of the membrane where at the boundary the membrane becomes flat, there is a continuous curvature change region between the cap and the boundary, which we call the “relaxation region” in the following derivation. The shape function of this relaxation region that detaches from the sphere will determine its bending energy. To briefly describe the solution, consider the shape function, where measures the height of the membrane and measures the azimuthal distance from the z-axis at the origin, given a membrane bending modulus. We make three simplifying assumptions: (1) Azimuthal symmetry: For any real function (2) Negligible spontaneous curvature: (3) Negligible membrane tension:, and therefore. We solve for the shape function under the following three boundary conditions: (1) Membrane is continuous at transition: ; (2) Membrane is differentiable at transition: ; (3) Membrane becomes flat at end of relaxation:, where is the radial distances from the z-axis to the end of the relaxation function.

The bending energy of the relaxation region *E*_B,relax_ then turns into:

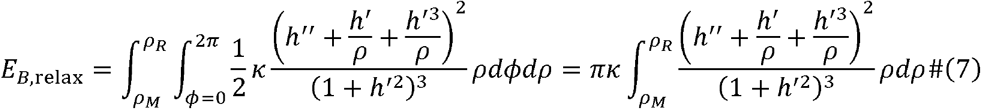

The shape function *h(ρ*) of the relaxation region that minimizes *E*_B,relax_ is then solved numerically (the detailed result is in SI section 3.3). To compare the prediction of the simple cap and relaxation model,

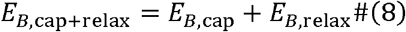

with our analytical results, we use *r*_*M*_ = *r*_G_ *l* _b_, to include the membrane displacement by the lattice and the harmonic bonds. We set *ρ*_*R*_=135nm, which is half of the boundary box size of the simulation.

This analytical function assumes that the tension is zero, and therefore the area energy does not contribute to the total energy, which is distinct from the energy function used to minimize the continuum membrane. To assess the impact of the additional area energy term on the predicted shape function of the relaxation region and its energy, we therefore also evaluated the minimum energy solution with the addition of the area energy (see SI). The solution for *h*(*ρ*) is no longer analytic due to the addition of the boundary conditions on the domain size. We solve for this iteratively (see SI), finding that this more accurate model predicts only relatively small changes in *h* (*ρ*) and a <2% change in the bending energy. We therefore conclude that deviations between theory and our continuum membrane simulations are not due to inconsistencies in the membrane tension assumptions.

### II. B Numerical Methods

#### II.B.1. Background on the Finite-element model for defining surface geometry

We use the Loop subdivision limit surface ^48^ to model the continuous and smooth membrane surface as described in previous research ^38^ and implemented in our previous work ^39^. This method uses a smooth limit surface to which the Loop subdivision algorithm converges when iteratively applied to a finite element mesh. The advantage of this method is that instead of measuring energy along the discrete and discontinuous triangular mesh, here referred to as the ‘control mesh’, the energy is evaluated on the smooth limit surface which has continuous first and second (excepting irregular vertices) derivatives needed for calculating forces and curvatures. Mathematically, any arbitrary point **p** on the limit surface *S* is determined by a weighted average of the surrounding backbone vertices ***q***_*l*_ of the control mesh (Fig 1C,D). The point ***p*** can be represented by barycentric coordinates(*u,v*)on its local element triangle, facilitating a spline fit using shape functions ^49^. We provide further implementation details in SI 3.1 and 3.2.

The energy function in Equation 1 is widely used in continuum models of membranes, and we verified that our numerical implementation correctly evaluates the energy defined above (Fig 1). In previous work we showed that our numerical implementation reproduces shape changes including with the addition of volume constraints for enclosed membranes (e.g. vesicles) ^78^. We further quantified shape and energy variations with local changes to spontaneous curvature as produced by helix-inserting proteins ^78^ and by scaffolding of the membrane by rigid clathrin lattices ^13^. We tested that the addition of the regularization energy *E*_reg_ to ensure the reshaping of the triangular mesh does not disrupt convergence to a low energy structure^78,79^. We note that we have not implemented edge flipping or dynamic remeshing of our triangular mesh^86,87^. As a result, we limited our lattices to a radius of up to 40 nm (max azimuthal radius is 59 nm, which is the spherical radius of Gag attached membrane used in the simulation). With larger lattices, the numerical reshaping of the membrane topology leads to large changes in mesh triangles that limit numerical convergence, and future work will extend the scale of shape changes accessible via the addition of dynamic remeshing.

#### II.B.2. Boundary Conditions for the continuum membrane model

All simulations are run under periodic boundary condition in **X** and but **Y** not in the **Z** direction, to mimic modeling a patch of a much larger membrane surface. For the periodic boundary, we create three rows of temporary or ‘ghost’ vertices outside the edge of the mesh with positions determined by translation of the mesh vertices at the opposite edge. Three rows are needed because with the limit surface set-up, each vertex requires 12 surrounding vertices to compute a position **p** on the limit surface. See SI section 3.2.4 for implementation details on the periodic boundary condition.

#### II.B. 3 Numerical integration of the continuum energy function across the surface

We use the second-order Gaussian quadrature scheme to numerically estimate the integral in our energy function. In our previous research, we demonstrated that the second-order Gaussian quadrature produces membrane structures with adequately accurate and converged estimations of the energy, while having a considerably lower computation time cost compared to integral methods with higher precision ^78^.

#### II.B.4. Self-assembled lattices from NERDSS simulations

We previously ran stochastic, particle-based reaction-diffusion simulations of Gag lattice self-assembly with the NERDSS software ^80^. The highly coarse-grained rigid structure of each Gag monomer and their interactions with one another were designed from the experimental Cryo-ET structure of the immature Gag lattice^6^. The model and methods were previously published ^16^, so we briefly summarize here. Each Gag monomer has 5 interaction sites for another Gag monomer, one homodimer site, two hexamer sites that bind to form a hexameric ring, and two trimer sites that form a trimeric ring. The monomers assemble into spherical shells with a tri-hexagonal lattice containing up to *N*_sphere_ =3700 monomers with a radius *r*_G_ = 50 nm. Because a rigid hexagonal lattice does not perfectly tile a sphere, the lattice also contains defects, which is consistent with the experimental structures ^16,81^. Depending on the initial conditions and strength of interactions, the Gag lattices do not always form complete lattices but instead become kinetically trapped in intermediates with varying sizes and morphologies. We sampled from these Gag lattice structures that are stochastically assembled during our simulations as a function of monomer numbers and the morphology. The morphology of lattices can be quantified by its compactness, or a regularity index that measure deviations from ideal spherical growth. Note that these solution simulations did not include any membrane. The relationship between the number of Gag monomers *n*_Gag_ and azimuthal radius *ρ*G of a compact or ideally spherical Gag lattice cap is given by: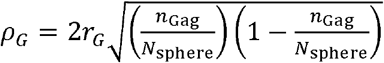.

#### II.B.5 Controlling membrane tension by specifying the equilibrium area

The total membrane energy depends on the area constraint energy *E*_*A*_, which is parameterized by *μ* _*A*_ and the equilibrium area *A*_0_. Experimentally, most living cells have a relatively low tension, in the range of values: typical membrane tension 0.01 pN/nm to 0.04 pN/nm ^82^ and in higher tension cellular membranes such as cytoskeleton-scaffolded membrane in moving cells and membranes of epithelial cells under osmotic stress are 0.15 pN/nm to 0.30 pN/nm ^83,84^. To keep our remodeled membranes at nonzero tension while minimizing *E*_tot_, we optimize the values of *A*_0_ (see Fig S3). Note that this is approach is different from assuming negligible tension in the system, i.e. *E*_*A*_ =0. However, the minimized surfaces do tend to have a quite low, but nonzero, *E*_*A*_. We do not use a zero-tension state, as this selects for nonphysical membranes with gradual curvature changes in the relaxation region that eliminate bending energy costs in this region (not at the cap). We verify that we can achieve a reasonable tension as well as we vary *A*_0_ (Fig S4). To imitate the scenario where lipids are allowed to diffuse freely in and out of the boundary, we run a series of simulations with changed spontaneous area and get the minimum total energy of all these simulations with a confidence interval calculated with bootstrap resampling test.

#### II.B.6 Definition of initial membrane shape under the protein lattice

In previous studies of membrane bending^78^, the membrane was initialized in a flat state, which was feasible because the equilibrium configuration remained close to flat, resulting in small forces. In our case, starting from a flat membrane configuration attached to the Gag lattice is challenging. It leads to large forces in the cap region and minimal forces near the end of and beyond the relaxation region. This uneven distribution of force together with a large maximum force complicates the energy minimization process: if we use the step size according to the empirical step-size rule, it becomes very small and slows convergence. In contrast, using a large time step results in mesh distortion, ultimately causing the FEM solver to fail. To resolve these difficulties, we initialized the system with a pre-curved configuration. The membrane is initially put in a near-to-equilibrium state that is based on the analytical solution to the membrane shape solved with no tension and the small gradient approximation. However, in this analytical solution, the membrane bending energy converges to zero at infinite distance and results in infinite membrane bulge height, which is not feasible (or physically realistic) within a finite bounded simulation. Therefore, we give the model a relaxation radius (*ρ*_*R*_) and calculate the lowest energy state analytical solution with that relaxation radius.

#### II.B.7. Computational optimization and energy minimization of the coupled system

We use nonlinear conjugate gradient method with Wolfe condition ^85^ and simple local gradient descent to propagate the membrane shape to minimize the total energy (see Fig S5). The direction of propagation of the Gag lattice is based on the force exerted by the membrane on the Gag lattice. The forces and their numerical implementation following from the bending and area energies have been defined in previous work ^78,79^, and the forces from the harmonic coupling to the protein lattice are defined in Eq. 4. The step size is controlled as a small value satisfying the empirical rule, max(‖force ‖) stepSize ≪lFace, where lFace is a constant hyperparameter (Table S3). There is a free variable here, which is the number of propagations of Gag per propagation step of the lipid membrane (see Table S3). In the simulation, *k*_hb_ is set to a small initial value 1.5 pN/nm and gradually increased to a final value of 10^3^ pN/nm, which makes sure that the finite element mesh does not deform or stuck in small propagation step size due to big initial force.

#### II.B.8. Theoretical lattices from Monte Carlo optimization

To establish a theoretical baseline to compare with the self-assembled lattices from NERDSS simulation, a Monte Carlo optimization program is used to generate theoretical Gag lattices with given lattice size and eccentricity, *e*. For an ellipse defined as:

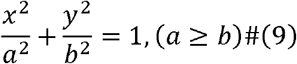

Where*a* is the major axis and *b* is the minor axis. The eccentricity *e*.is then defined as:

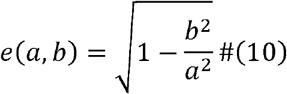

Here, the Gag monomers are simplified to nodes lying on the spherical surface, fully ignoring the orientation of interaction. Specifically, the Monte Carlo optimization starts with a set of randomly placed nodes representing the Gag monomers on the Gag spherical surface within boundary. Even if the Gag monomers are simplified to nodes lying on the spherical surface, fully ignoring the orientation of interaction, generating an “ideal” mesh that has “evenly placed nodes” that are *l*_gg_ =4.7 nm (average Gag-Gag interaction length) apart within a particular geometric boundary can be challenging, due to the topological layout of the nodes rely strongly on the shape of the surface. To generate an ideal mesh that has evenly placed nodes, we used a Monte-Carlo simulation to start with a given number of nodes, and let the nodes propagate on the surface based on Lennard-Jones Potential, with the minimum Lennard-Jones energy at where nodes are exactly *l*_gg_ apart. To accelerate the simulation, we apply simply the summed Lennard-Jones potential of the system to only calculate the potential between each node and their closest neighbor using KDTree. The method is further elaborated in SI 3.4.

## III.Results

### III. Stochastically assembled lattices vary in their sizes and regularities, leading to varied membrane energy

As we show in Figure 3, when we bend the membrane via lattices that were stochastically assembled, the total membrane energy cost is higher when the lattice contains a larger number of Gag monomers. An increase in membrane energy cost is consistent with theoretical predictions for spherical caps of increasing size (Eq. 6), primarily driving by the bending energy (Fig 3b). However, the value of the membrane energy from the stochastic lattices is frequently much higher than that predicted by theory, even with the finite-size boundary correction (Fig 3b). The simple cap theory assumes the relaxation area energy is zero and thus disregards any boundary size effect. The cap and relaxation theory accounts for the boundary size effect (Eq. 8, Methods), but this is still insufficient to explain the discrepancy with the simulations. Visual inspection of the Gag lattice structures suggests that more irregular or noncompact lattices result in larger membrane energy and thus larger deviations from the theoretical predictions. For example, the 227-Gag configuration is highly irregular relative to an ideal spherical cap and shows a significantly higher energy than predicted, whereas the 157-Gag case is more regular and only deviates moderately. Interestingly, despite the irregularity of the lattices, the membrane also prefers to bend into a cap-like deformation, with minimal ‘wrapping’ around the irregular perimeter (Fig 3c). As we show below, this observation is confirmed by systematically varying the regularity of the lattice by using idealized meshes that deviate from perfect sphericity, as quantified by their eccentricity,*e*.

**Figure 3.**
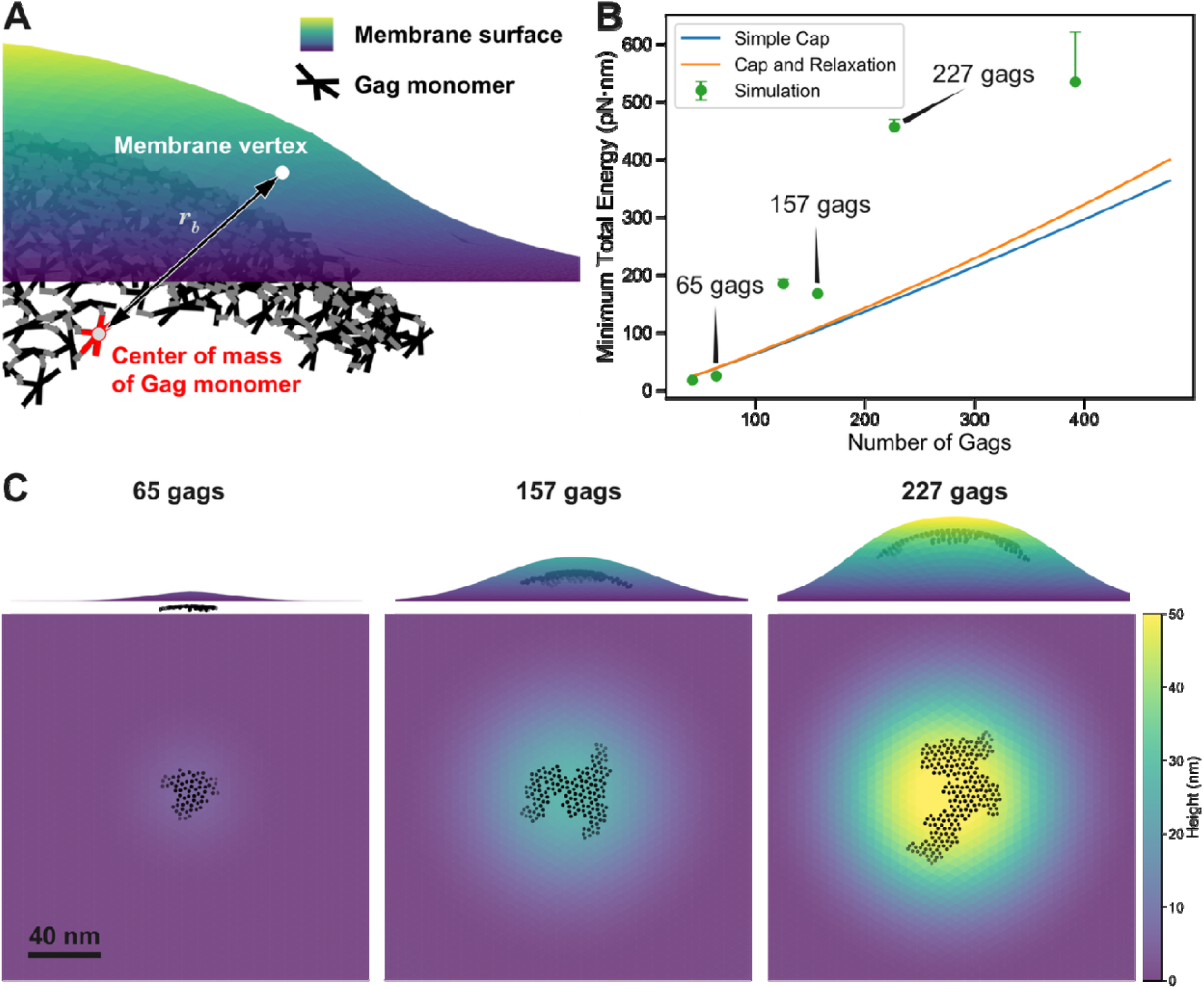
Energy increases with the size and irregularity of the Gag lattices. A) Rigid lattices are connected to the membrane surface via harmonic bonds. Gag monomers are in black, surface is contour in colors by height. The average surface area per Gag in a complete lattice is 11.82 nm^2^. B) The minimized total membrane energy measured for the continuum membrane surface plus the harmonic bonds (Methods). Gag lattices are stochastically assembled from NERDSS simulations. Theoretical curves are calculated based on a simple (discontinuous) cap model (blue curve, Eq. 6), and accounting for a continuous curvature relaxation with the boundary size effect (orange curve, Eq. 8). Error bar represents 95% confidence interval from bootstrap resampling. C) Energy-minimized structures of Gag lattices with the continuum membrane surface show increased membrane remodeling around larger lattices. A membrane energy minimization is run with bending modulus of 83.4 pN·nm, area stretching constant of *µ*_*A*_ =250.0 pN/nm, and 270 nm × 270 nm boundary, which is maintained in all simulations.

### III. B. Membrane bending energy is higher for more eccentric lattices

To test the role of lattice asymmetry on the membrane bending energy, we constructed artificial meshes that represent the centers-of-mass of each Gag monomer using Monte Carlo sampling (Methods). We systematically varied the eccentricity of these lattice caps, keeping the lattice curvature constant at *r*_*G*_^-1^ =(50*nm*)^-1^ (Fig 4a). Higher eccentricity reflects an increasingly elongated arrangement of the lattice “monomers”. Our simulation results show that the higher the eccentricity of the lattice, given the same number of Gag monomers, the higher the total membrane energy, >99% of which arises from the bending energy (Fig 4b, Fig S6). The minimal contribution of the area stretching energy to the total energy is expected, as we are focused on a regime of low membrane tension. The more eccentric lattices clearly drive a larger invagination of the membrane given the same number of monomers (Fig4c, Fig S7). Thus we see a deformation with a similar height when 137 monomers bend the membrane with *e* = =0.87 vs 302 Gags at *e*=0 (Fig 4). The height of the deformation, measured simply as the difference between the maximum and minimum z-coordinate of our control mesh vertices (Fig 4c), correlates well with the energy (Fig 4b), as expected from theory of spherical caps. Here again we see that despite the non-circular perimeter of the eccentric lattices, the membrane still bends into a relatively circular or radially symmetric invagination.

**Figure 4.**
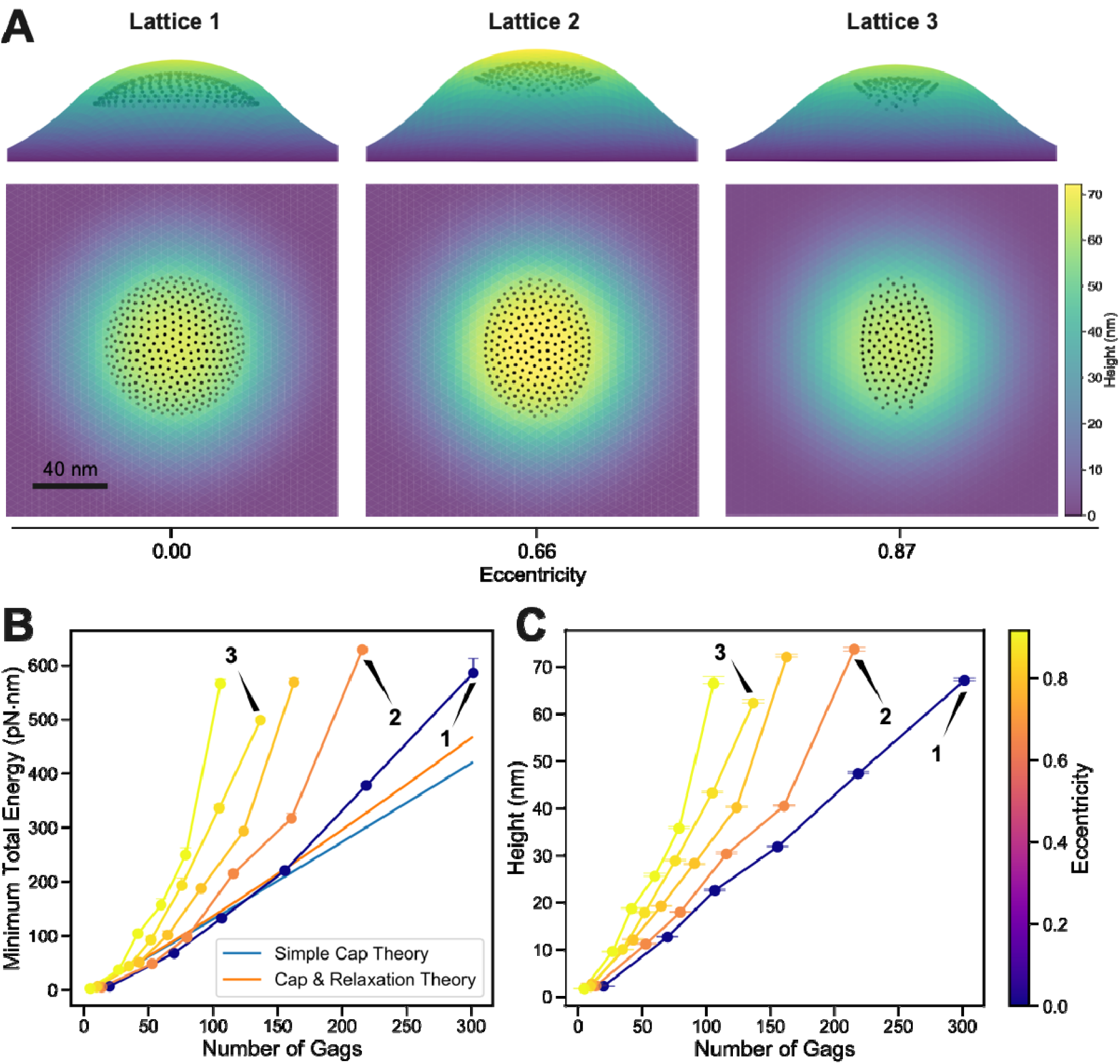
Membrane energy cost increases with the eccentricity of the lattice. A) Idealized lattice structures are attached to the membrane and energy-minimized. From left to right the size of the major axis remains the same at 40 nm, but the eccentricity increase. Left has 302 Gags, center has 216 Gags, and right has 137 Gags, illustrating how a similar sized invagination is produced by a small number of Gags when they are highly eccentrically arranged. B) The minimized total membrane energy when attached to Gag lattice of different size and eccentricity assembled with Monte-Carlo method versus the number of Gags in the lattice. Error bar represents 95% confidence interval from bootstrap resampling. Over 99% of the total energy is due to the bending energy (Fig S6). C) Membrane cap height increases as with the eccentricity of the lattice with minimum variance. The height of the membrane cap at minimized total membrane energy when attached to Gag lattice of different size and eccentricity assembled with Monte-Carlo method versus the number of Gags in the lattice. Error bar represents 90% bilateral confidence interval from joint bootstrap resampling. The height of the invagination and the energy are closely correlated.

The penalty associated with bending the membrane with more eccentric lattices is not linear in the size of the lattice, as suggested by the energy growth in Fig 4. For small lattices, the cost of eccentricity is modest, whereas for large lattices, the cost of eccentricity grows rapidly (Fig 5). This observation further confirms our previous hypothesis: more irregular or eccentric lattice configurations impose greater energetic penalties on the membrane, and this is more visible for larger lattices.

**Figure 5.**
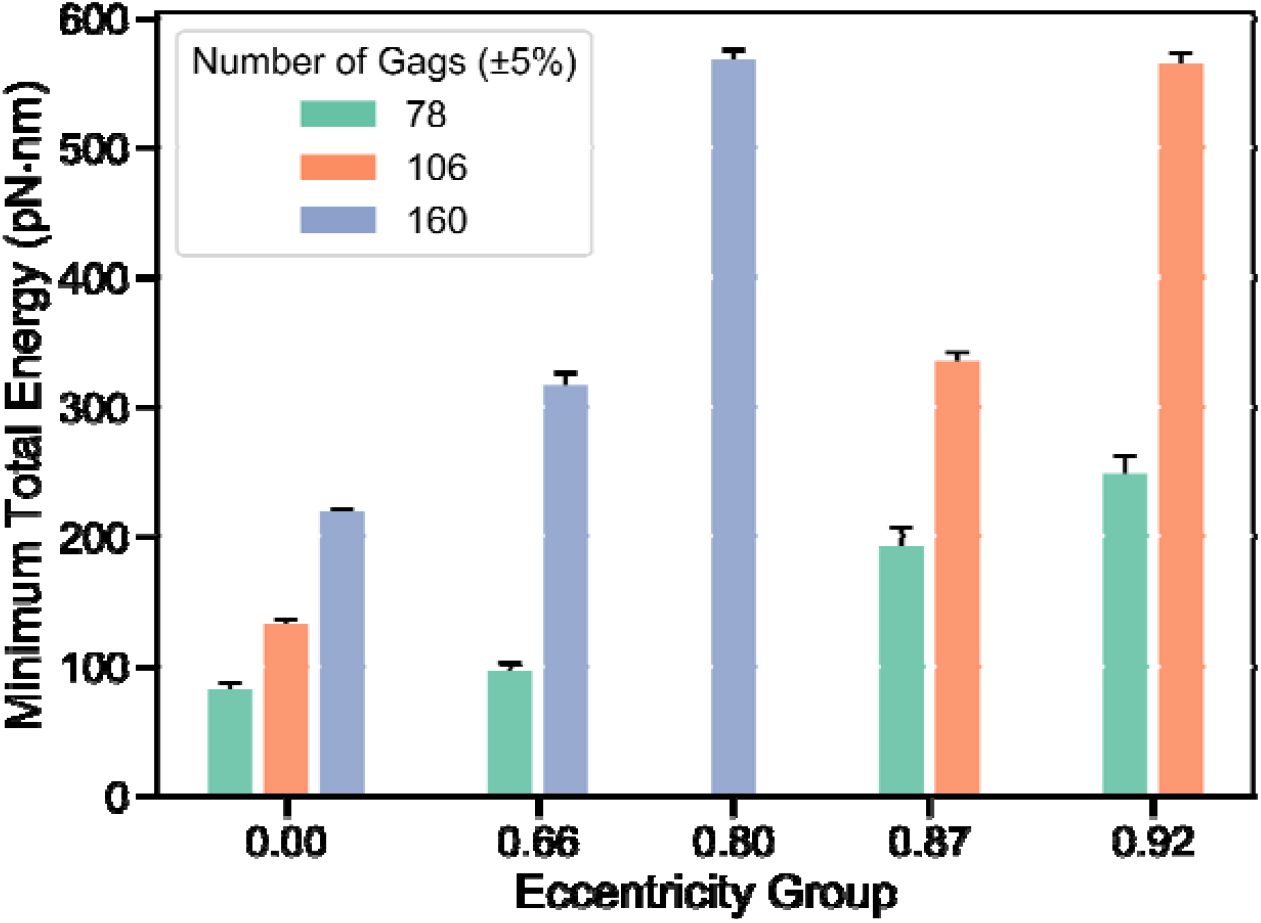
The membrane bending energy cost with increasing eccentricity rises more rapidly for larger lattices. For smaller lattices (green), increases in eccentricity have modest impact on the bending energy. For larger lattices (orange and blue), the energy grows more rapidly with eccentricity. The minimized bending energy of the membrane when attached to Gag lattice assembled with Monte-Carlo method versus the eccentricities, grouped by the number of Gags with a ±5% tolerance in each group. Error bar represents 95% confidence interval from bootstrap resampling. First, a script is run to scan through all integer values from between the minimum and maximum number of Gags in the dataset and identifies the simulation trials whose number of Gags values fall within ±5% of the target number of Gags. If at least three such trials are found, their indices are recorded and plotted here.

We note that lattices with *e*=0 form effectively ideal spherical caps, and therefore we expect our numerical simulation results to agree well with the theory in Eqs. 6 and 8 plotted in Fig 4b. While the agreement is quite strong, the growth rate is somewhat faster for the continuum model. For the small lattices, the continuum membranes have a *lower* energy than theory predicts, whereas for the larger lattices, the continuum membranes have a *higher* energy than theory. At the largest structure, the continuum membrane predicts a higher energy by ∼20%, which cannot be explained by theoretical assumptions on the role of membrane tension, for example (see Methods and SI). By comparing the shape functions of the membrane invaginations predicted by theory with the corresponding cross-section of our minimized continuum model solutions, we see they do not overlap perfectly (Fig S8). These deviations likely stem from the assumptions of the theory, which predicts a complete cap that is then continuously connected to the relaxation region. In contrast, the continuum model can simultaneously modify the shape of both the cap and relaxation regions. For small lattices we see that the continuum model seems to find a smaller radial cap deformation due to the offset of the rigid Gag lattice below the membrane via its harmonic bonds, thus lowering the bending energy (Fig S8). For the larger lattices, the opposite seems to occur, with a larger cap region before relaxation to the flat membrane, thus increasing the bending energy (Fig S8). Because we are performing energy minimization, our solution could also represent a local minimum, based on the initial height we applied to the membrane and the limits of the periodic boundary conditions. Increasing the membrane size or changing the initial starting configuration to match the theory could result in tighter agreement between both shape and energy.

### III.C. Low global tension in the system

We verified that in our energy-minimized systems of coupled lattices and membranes; the tension did not vary significantly (Fig 6). By calculating membrane tension using Equation 3, we showed that our membrane tension is controlled within physiologic regimes (or lower). Although the tension does increase systematically with the size of the lattice and with its eccentricity, the tension remains below 0.014 pN/nm, with estimated upper bounds on most cell types reaching 0.04 pN/nm ^82^. We nonetheless validated that this growth in tension is not the reason for the increasing total energy (Fig S9). The area constraint energy, which relates to the membrane tensions (Eq 3), contributes minimally to the total energy of the optimal structures () (Fig 6b).

**Figure 6.**
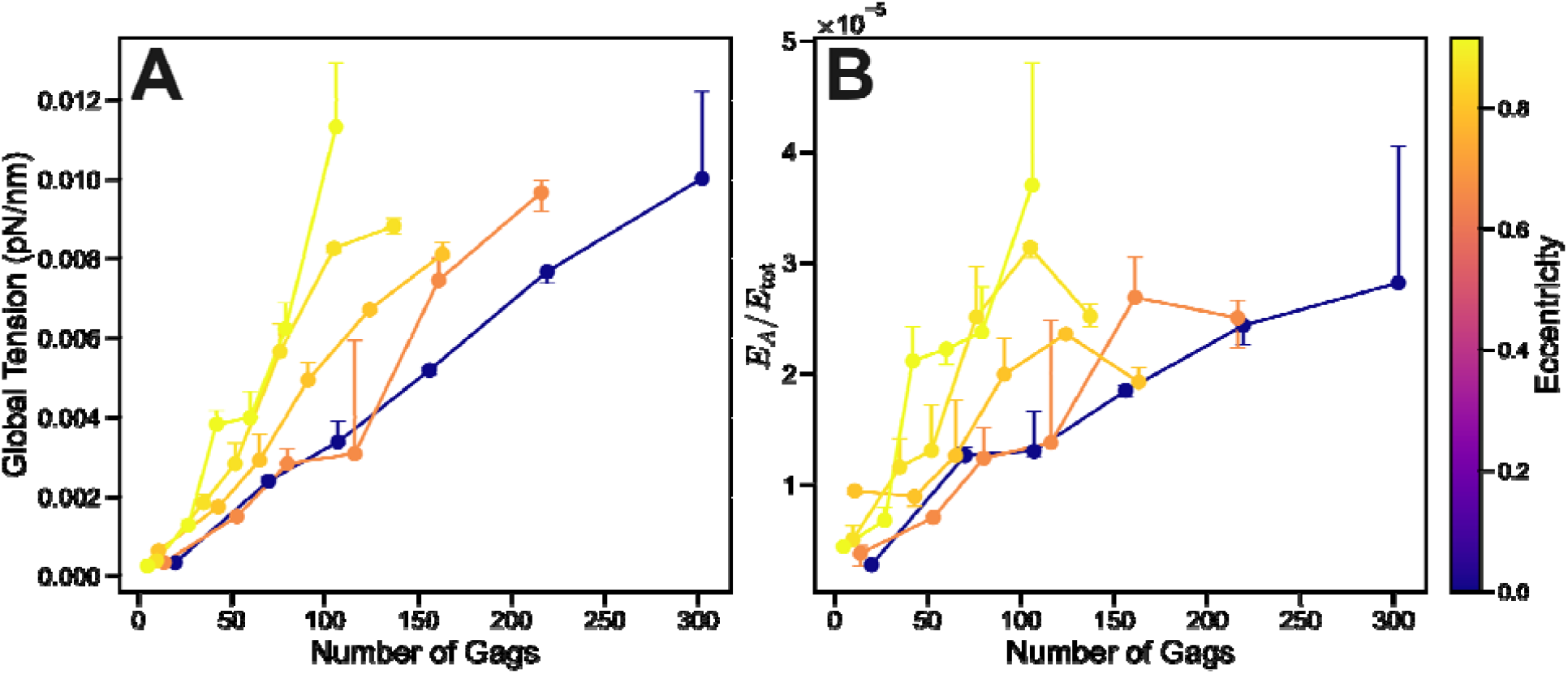
Our membrane tension is constrained to low (physiologic) values that contribute minimally to total energy. A) Global tension of the membrane when attached to ‘Gag’ lattices assembled with Monte-Carlo method was measured using Eq. 3 (Methods). Although tension does increase with number of Gags and eccentricity (color bar), it remains within or below physiologic regimes of 0.01-0.04 pN/nm. Same simulation data used in Fig 4 and 5. B) The fraction of total energy coming from the area constraint is very small for all lattices, <5×10^−5^. Error bars represent 90% bilateral confidence interval from joint bootstrap resampling.

### III.D. Density of monomer attachments to the membrane has a minor impact on bending energy

To test whether the density of monomer attachments to the membrane impacted the bending and total energy, we increased the separation *d*_*MM*_ between the monomers while keeping the perimeter size fixed by an azimuthal radius of 30nm (Fig 7a). With fewer attachments to the membrane, we do see that the invagination by the lattice becomes slightly higher (Fig 7a). However, this has negligible impact on the bending energy for the higher densities, where from *d*_*MM*_ = 3.76 to 5.64*nm*, no significant change occurs in bending energy (Fig 7b) or total energy (Fig 7c). For reference, we include the simple theoretical cap estimate of the bending energy (Eq 6), which is relatively close to the simulated membrane bending energies (Fig 7). The simulated values must exceed this theoretical minimum because of the addition of tension and boundary effects. For the lower density lattices, *d*_*MM*_ = 7.05 to 9.4*nm*, we do see a modest increase in bending energy that is statistically significant, indicating that the very sparse connections do drive higher invaginations which eventually creates a higher bending energy cost. However, the absolute differences are small — all under 14.2 pN·nm, which is less than 7% of the total energy. These results show that while the density of attachments between the rigid lattice and the membrane can increase the height of the deformation with lower attachments, its impact on membrane energy is minimal. Hence the size of the perimeter of the attached lattice is driving the bending energy costs.

**Figure 7.**
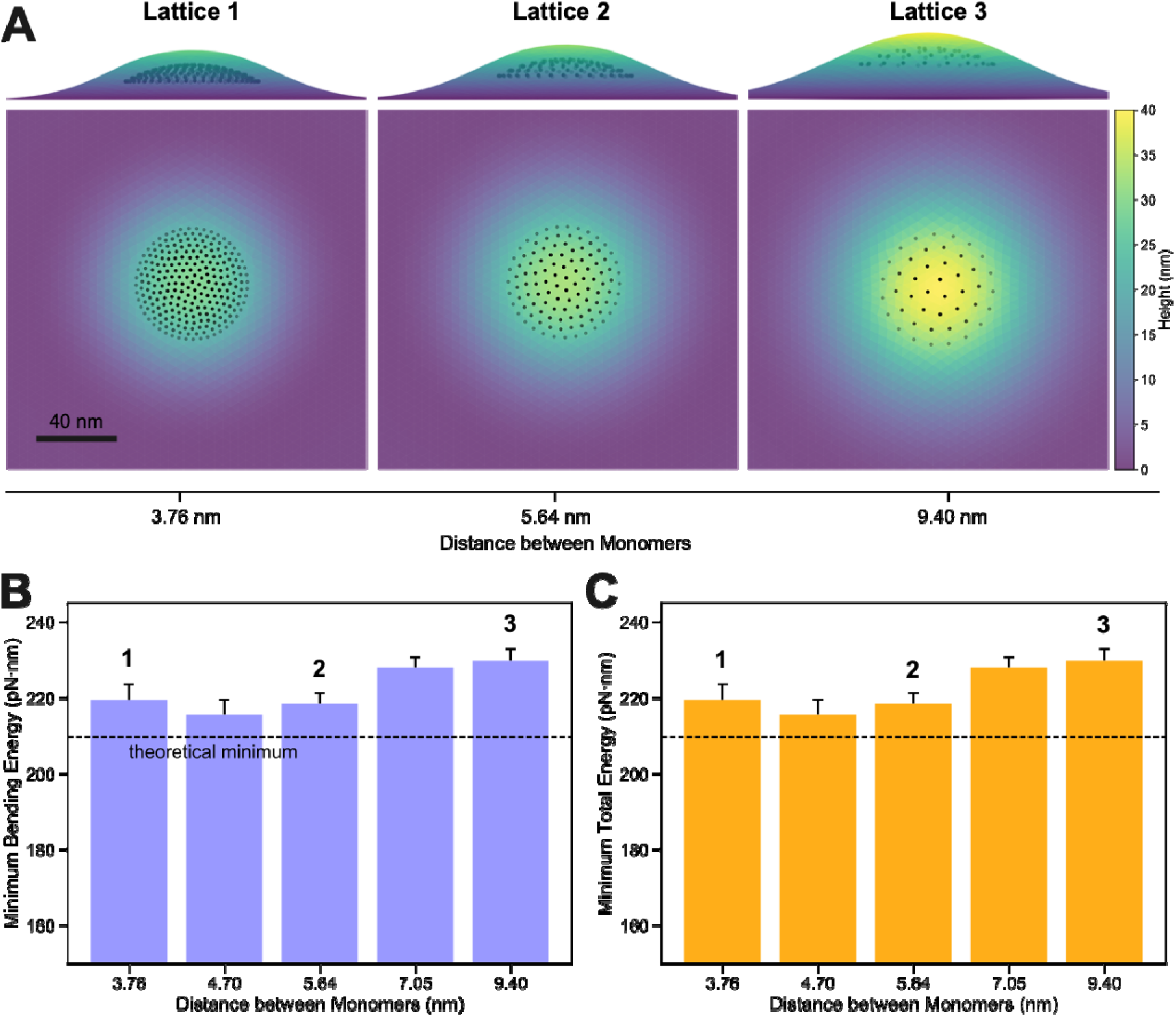
As density of Gags connecting to membrane is lowered, energy is not substantially changed. A) After fixing the size (azimuthal radius of 30 nm) and eccentricity () of the Gag lattice cap, we vary the number of equally spaced monomers/lattice points that attach to the membrane via the distance between monomers, from dense (to sparse (. The heights of the invaginations is similar (colorbar), with lattice 3 slightly higher. B) The bending energy is relatively constant as increases and fewer monomers attach to the lattice, indicating that it is the perimeter size (azimuthal radius) of the lattice that predominantly determines the bending energy. C) As expected, the total energy displays the same trends as the bending energy. Error bars show 95% confidence interval of minimum energy from bootstrap resampling. Theoretical minimum bending energy (209.6 pN⋅nm) assumes a simple cap model with no membrane tension (Eq. 6). We note there are statistically significant differences between the energy of the smaller lattices (*d*_*MM*_ < 5.6) relative to the *d*_*MM*_ =7.05 and 9.4nm lattices, indicating a somewhat higher bending energy once attachments are sufficiently sparse.

### III.E. Circumscribing cylindrical sector estimates the attached membrane energy

From the above results, we find the shape of the perimeter rather than the density of Gags is the major determinant of the attached membrane energy. Therefore, an assumption is that a Gag lattice with the same circumscribing cylindrical sector that defines the perimeter (Fig 8) leads to the same minimum bending and total energy of the attached membrane. This provides a consistent way to compare lattices of varying size and shape, regardless of Gag density (Fig 7) and encompassing all eccentricities. We indeed find that the minimized bending energies of membranes of all our lattices agrees well with the theoretical approximation when we plot it according to the cylindrical radius (Fig 8). The theory calculates the bending energy of a spherical invagination with the corresponding radius (Eq. 6-8). We observe that the agreement is not perfect, and indeed, the more spherical lattices (*e* − >0) tend to have energies that exceed the theoretical estimate. For the higher eccentricities, we see that the energy is typically lower, indicating that the bending energy cost is not quite as bad as one would expect for an eccentric lattice with major axis b vs a fully spherical lattice with radius b. This suggests that there is some radial asymmetry in the bending around the lattice of highly eccentric lattices that does reduce the surface area of the bending membrane. Finally, for the larger invaginations, they all tend to produce large bending energy costs than theoretically predicted (Fig 8a, r=40nm), as discussed in section III.B above, this is likely because of the variations in the minimized shape around the rigid lattice compared to the theoretical predictions.

**Figure 8.**
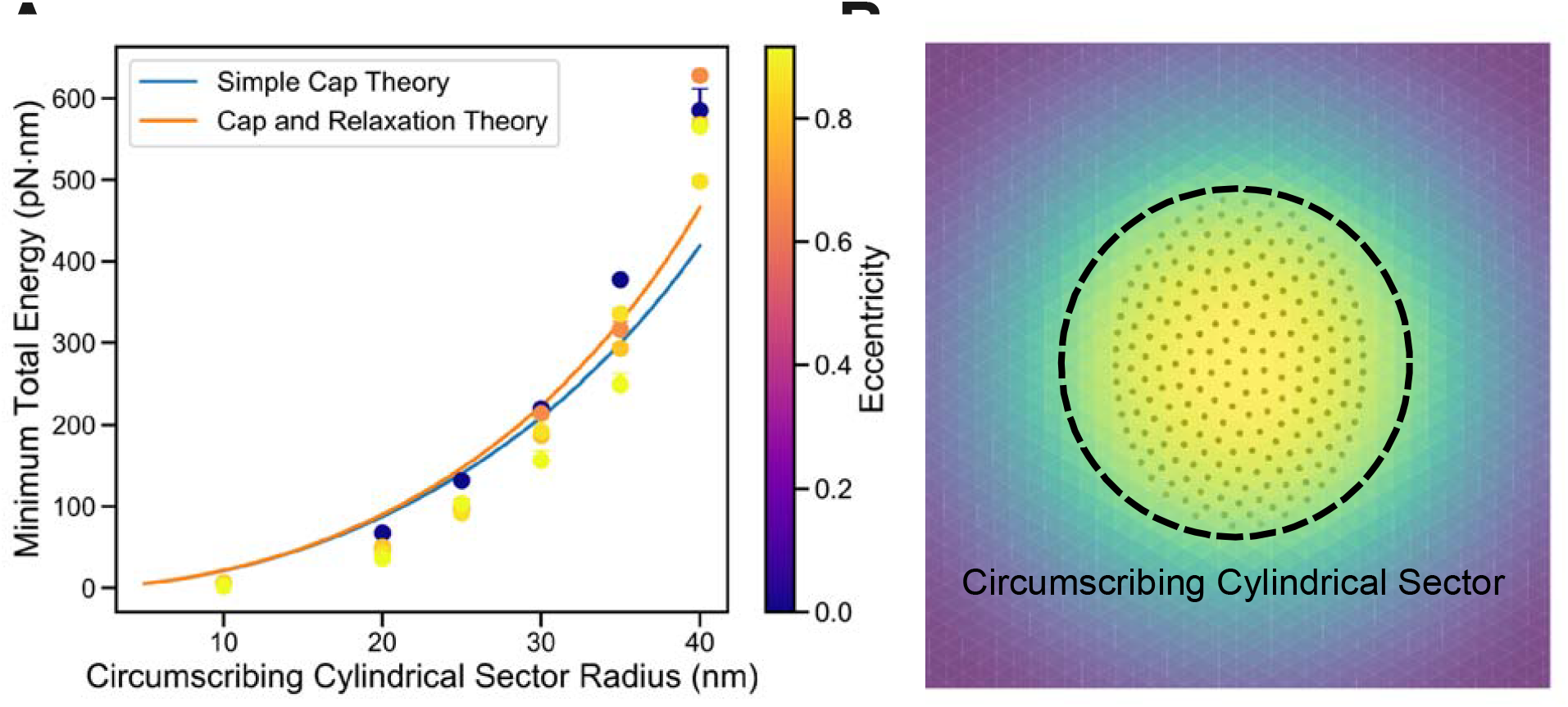
The membrane energy for all eccentricities is largely described by the radius of the circumscribing cylinder. A) Our data collapses onto a universal curve with relatively good agreement when plotted vs the radius of the circumscribing sphere rather than the number of Gags within the lattice. Same data points as previous figure, but with x-axis redefined. Simple cap theory from Eq. 6 (blue) and addition of boundary effects and relaxation without the small-gradient approximation from Eq. 8 (orange). B) We illustrate the circumscribing cylindrical sector here for a representative *e* = 0.66. It is the smallest cylindrical radii that fully encloses the lattice.

## IV. DISCUSSION

Our results clearly demonstrate that the membrane energy due to bending around a lattice scaffold increases not only with lattice size but with eccentricity of the lattice, despite containing the same lattice surface area. Although increases in eccentricity create asymmetry or irregularity in the lattice, the membrane still prefers to bend into a nearly radially symmetric invagination to minimize total bending energy. As a result, we can predict relatively accurately what the minimal membrane energy cost will be for any lattice regardless of its eccentricity using analytical theory, by calculating the radius of the cylinder that fully circumscribes the lattice. The discrepancies here between our approximate theory prediction and the continuum membrane predictions indicate that eccentric lattices induce some radial asymmetry into the cap-like invagination to lower its energy (Fig 8). However, compared to an ideal cap of the same surface area (e.g. monomer number), the increase in bending energy cost overwhelms this smaller benefit (Fig 4). Our findings emphasize that membrane deformation due to protein scaffolding is sensitive to lattice perimeter geometry much more so than the density of contacts between the lattice and membrane, suggesting that processes involving assemblies of irregular or non-axisymmetric protein on the membrane experience greater bending energy penalties given the same area of the deforming lattice. Overall, the dramatic increase in bending energy costs associated with asymmetric or irregular self-assembly pathways on the membrane would strongly select against their occurrence, despite being stochastically accessible in solution.

Our model framework is the first to systematically characterize how the eccentricity of spherically curved, rigid lattices pays a steep price in bending energy costs for all but the smallest structures. A limitation of this study is that the lattices were assumed perfectly rigid, including their links to the membrane. While protein lattice scaffolds are measured with bending moduli that are 10-fold or more higher than the soft membrane, the membrane could still induce compensating changes to the lattice to lower the total system energy cost. This may be particularly relevant for lattices with lower densities of membrane contacts—here we found that these lattices induced nearly the same bending energy cost, but because proteins are inherently flexible, a large separation between contacts may render those lattices less rigid. As the number of interactions per area of Gags increases, the lattice may become stiffer and less able to bend according to membrane shape, indirectly resulting in higher bending energy ^89,90^. To account for this, future models could implement an elastic energy term for the lattice itself, such as by treating it as a spring network or elastic limit surface ^75,77,91^. The mechanical properties of the lattices can be critical for their ability to not only deform membranes but to deform and withstand forces externally during processes like nuclear translocation of the HIV capsid^92^. The HIV lattice also grows and locks in defects due to the hexagonal lattice shape, locally lowering the density of Gag contacts to the membrane. Locking these defects in rapidly would help prevent the persistence of an irregular perimeter during viral scaffold assembly.

An important future direction will be the characterization of the role of local changes in spontaneous curvature, *C*_0_. Changes to spontaneous curvature can result from protein insertion to a single leaflet of the membrane, and for the Gag lattice, the insertion of the myristol group may bias the membrane towards negative curvature, helping to favor budding. Previous studies^88^ have demonstrated that spontaneous curvature can promote or suppress protein aggregation depending on the difference in the curvature of protein assembly and the spontaneous curvature of the membrane. Protein-induced curvature can drive membrane reshaping and tubulation even in the absence of motor proteins ^15^ and extending this framework to include stochastically assembled lattices could provide insight into how the mechanical membrane energy contributes to processes such as membrane scission and vesicle formation. Because of the importance of the perimeter, defects within the lattice would interfere with growth, although once they are embedded in the neighboring lattice, they would not increase the cost significantly based on our study of attachment density.

Given that thermal fluctuations penalize bigger membrane area and can assist in overcoming energy barriers, implementing Brownian or Langevin dynamics simulations with shape updates would allow exploration of equilibrium distribution or kinetic trajectories towards equilibrium evolving over time, beyond the static structures studied here. We effectively assume a zero-temperature membrane here, whereas shape fluctuations may reduce the energy cost of small irregularities during stochastic growth. Nonetheless, we would predict that a dynamically coupled self-assembly and remodeling simulation would eliminate proteins that bound to the edge of the lattice and substantially extended its eccentricity—the membrane bending cost would effectively increase the dissociation rate of these types of events much higher than they experience in solution. This approach will provide an open-source resource to the community for detailed understanding of membrane remodeling in cell biology.

## ACKNOWLEDGEMENTS

MEJ gratefully acknowledges funding support from an NIH R35GM133644. We thank Dr. Yiben Fu, Dr. Samuel Foley and Dr. Alex Sodt for helpful discussions.

