## Supplemental Information for "Membrane bending energy selects for compact growth of protein assemblies"

### Table of Contents

**1. Supporting Figures**......4

*1.1. Detailed results from simulation*...... ***Error! Bookmark not defined.***

*1.2. Geometry of the protein-scaffolded membrane*...... ***Error! Bookmark not defined.***

*1.3. Optimization flow chart*...... ***Error! Bookmark not defined.***

#### 1. Supporting Figures

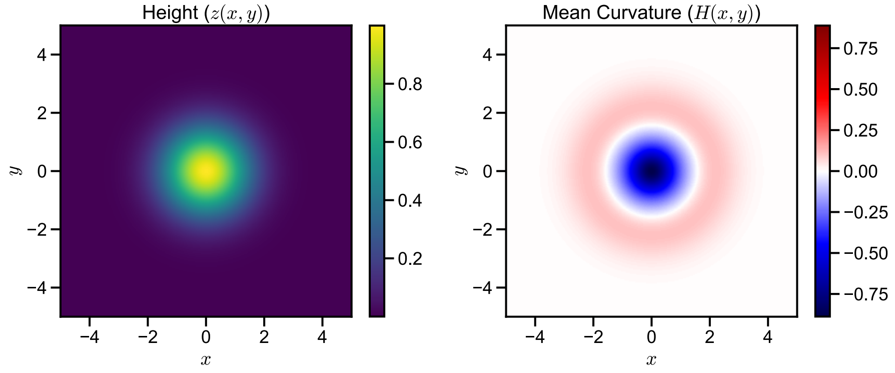

**Figure. S1. Heatmaps for height and mean curvature of a membrane bulge.** We see on the right panel that for a membrane bulge, there is a ring with near-zero mean curvature. This is because in this region, curvature is positive in the azimuthal direction and negative in the radial direction and thus cancelling each other out.

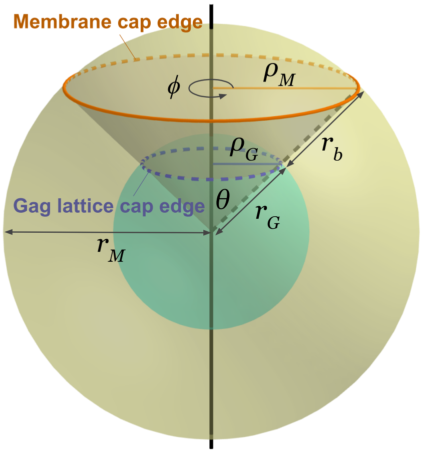

**Figure. S2. Graphical representation of the analytical model for the membrane and Gag lattice cap interaction.** The Gag lattice cap and the membrane cap are assumed to be concentric spherical caps with radius $r_G, r_M$ respectively.

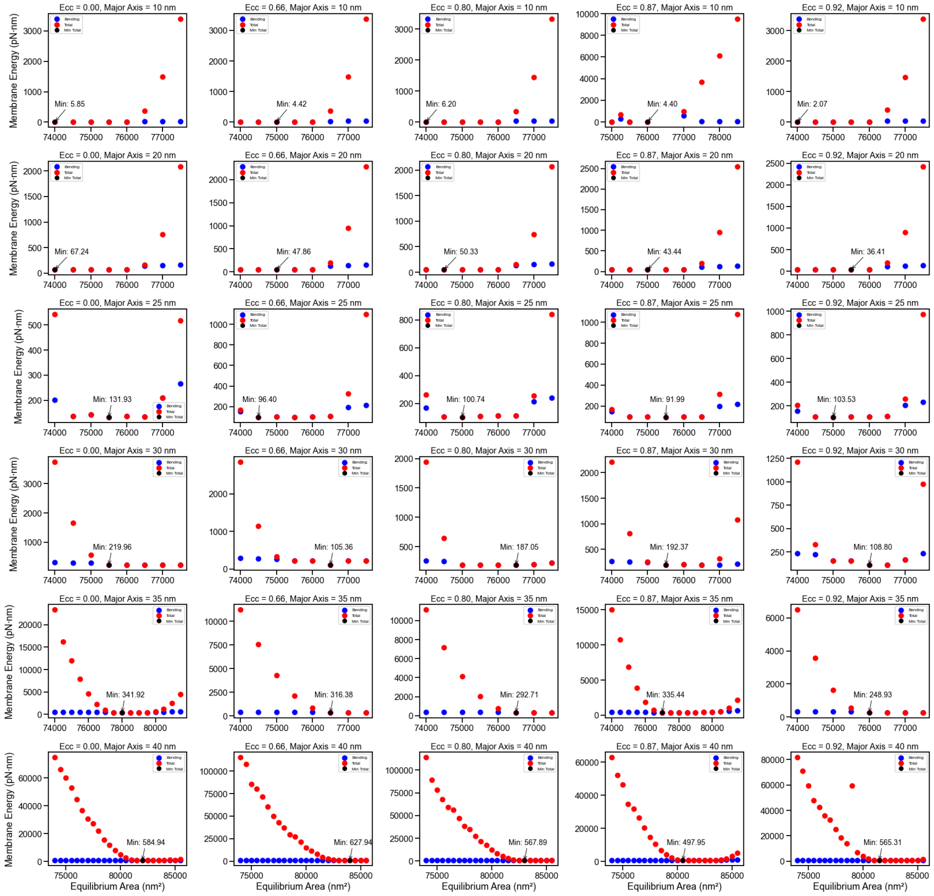

**Figure. S3.** Total energy and bending energy versus equilibrium area varying eccentricity and cap size. Each trajectory is shown as a separate data point here.

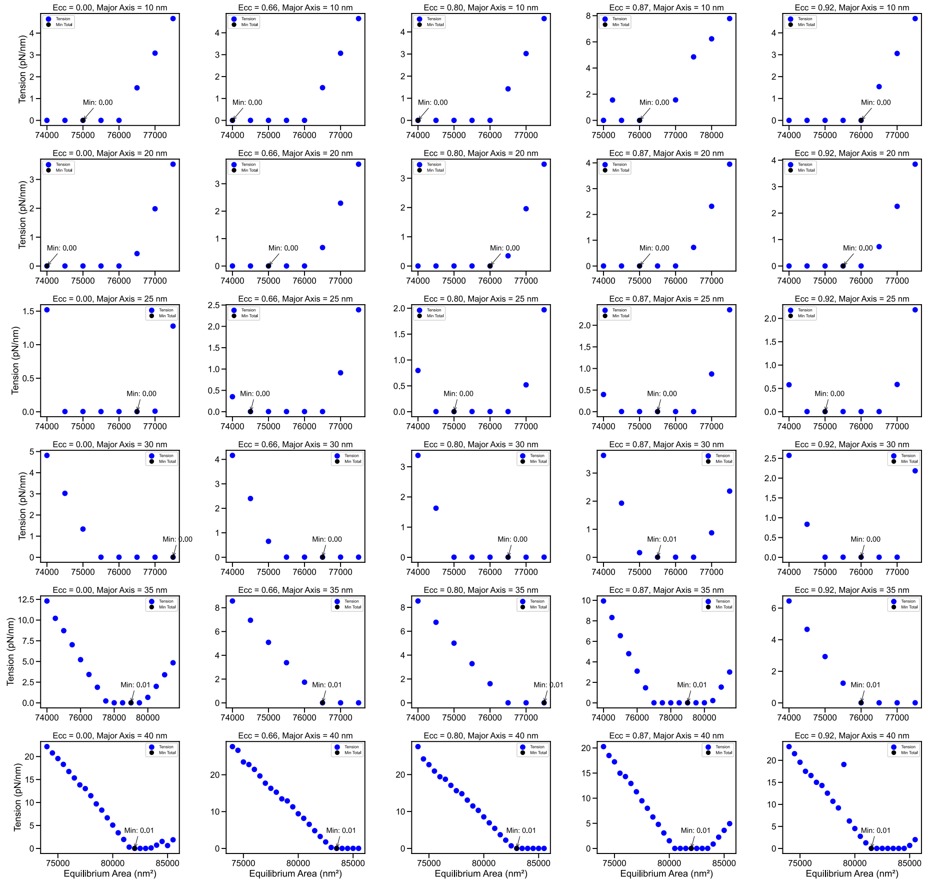

**Figure. S4.** Tension versus equilibrium area varying eccentricity and cap size. Each trajectory is shown as a separate data point here.

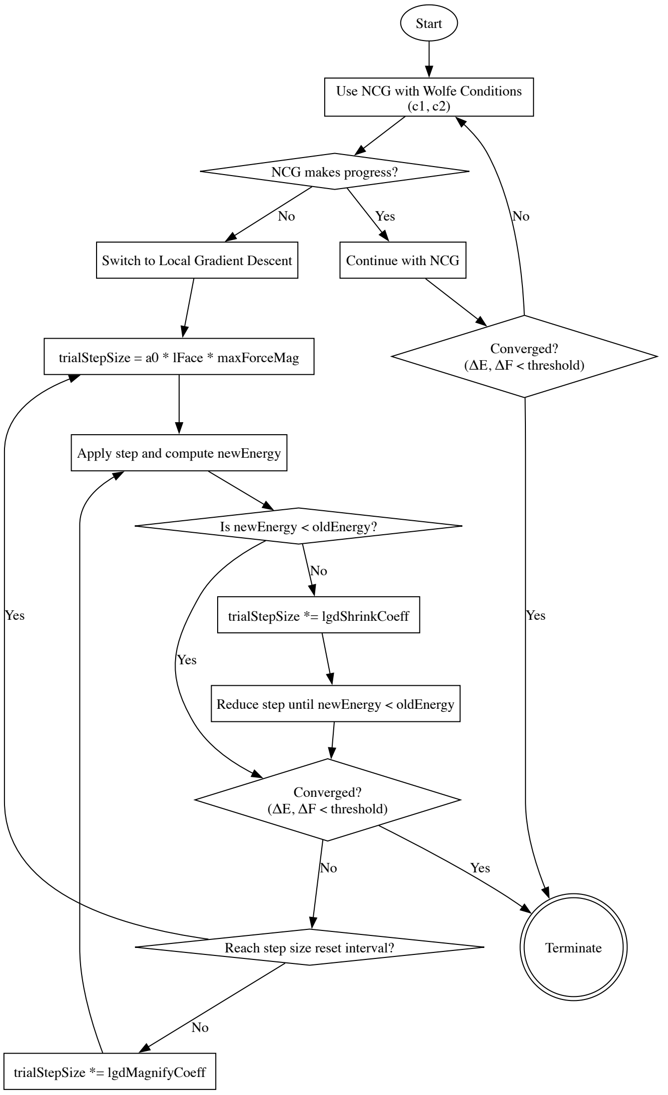

**Fig. S5. Flow chart for our numerical optimization**. We start with nonlinear conjugate gradient (NCG) with Wolfe condition and switch to a simple local gradient descent with adaptive step size if NCG fails. See Table. S3 for hyperparameters in the simulation.

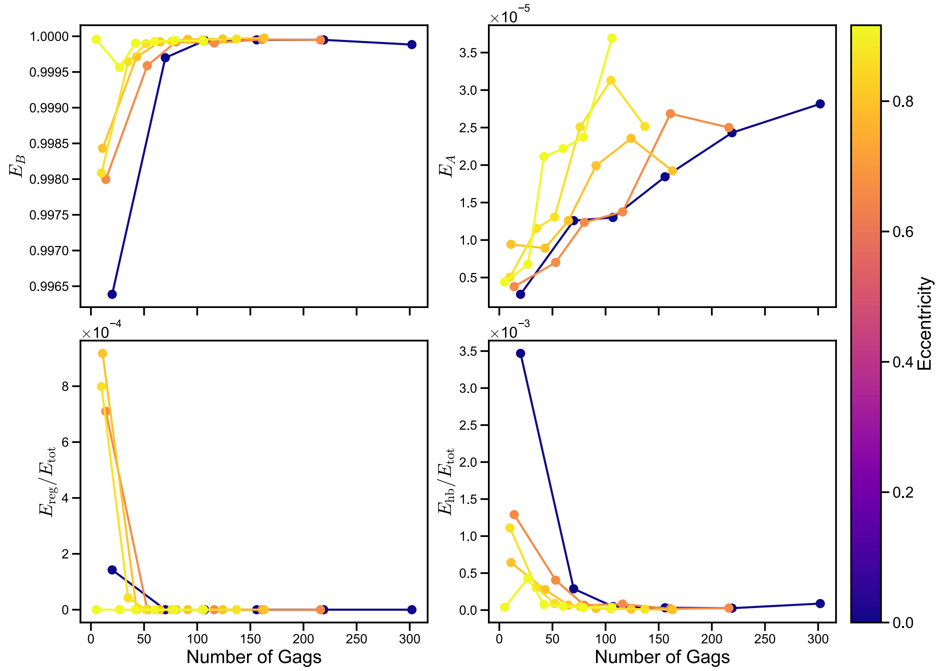

**Figure. S6. All energy terms other than bending energy contributes minimally to total energy.** Ratio of all energy terms (bending, area constraint, regularization and harmonic bond) to total energy of the membrane when attached to Gag lattice assembled with Monte-Carlo method versus the number of Gags with various eccentricities.

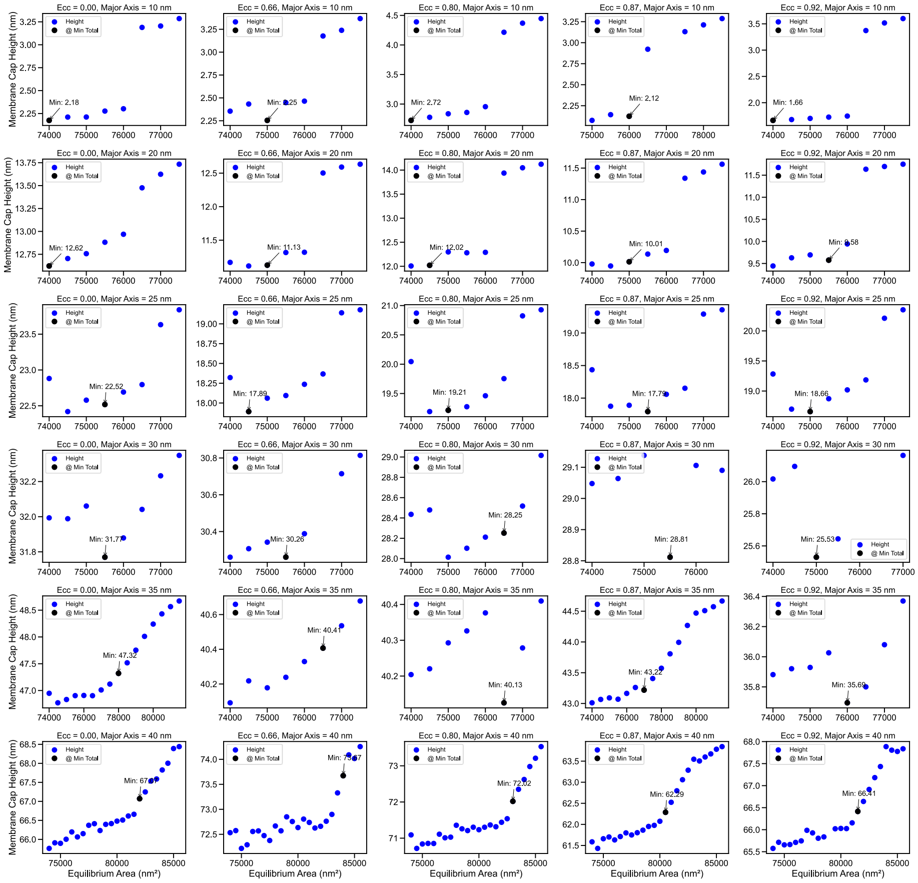

**Figure. S7.** Membrane cap height versus equilibrium area varying eccentricity and cap size. Each trajectory is shown as a separate data point here. Each row has a fixed Major Axis but an increasing eccentricity across the columns. This means that from the left column to the right column, the number of Gag monomers is decreasing.

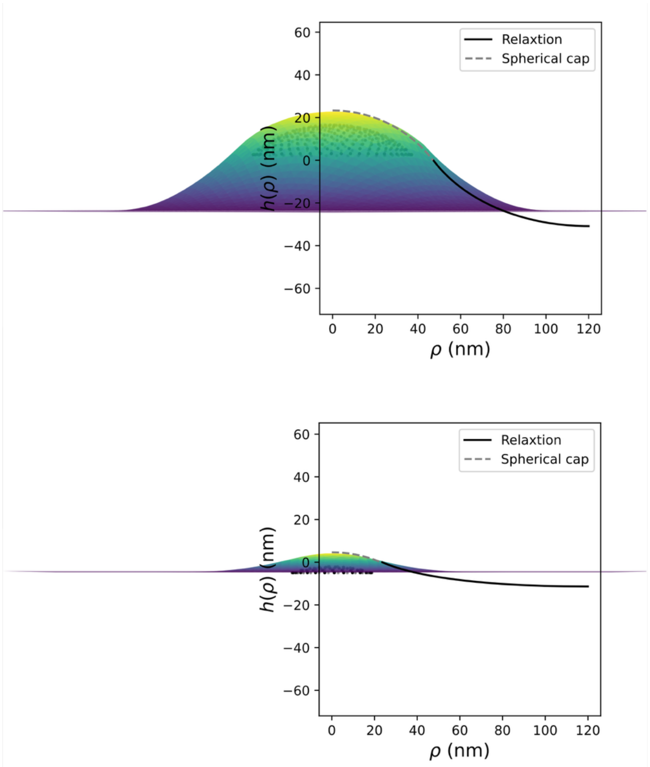

**Figure. S8.** Comparison between the membrane the theoretical and the simulation shape profile $h(\rho)$. Top is for a radius of 40nm, and the bottom is 20nm (radius of the Gag lattice).

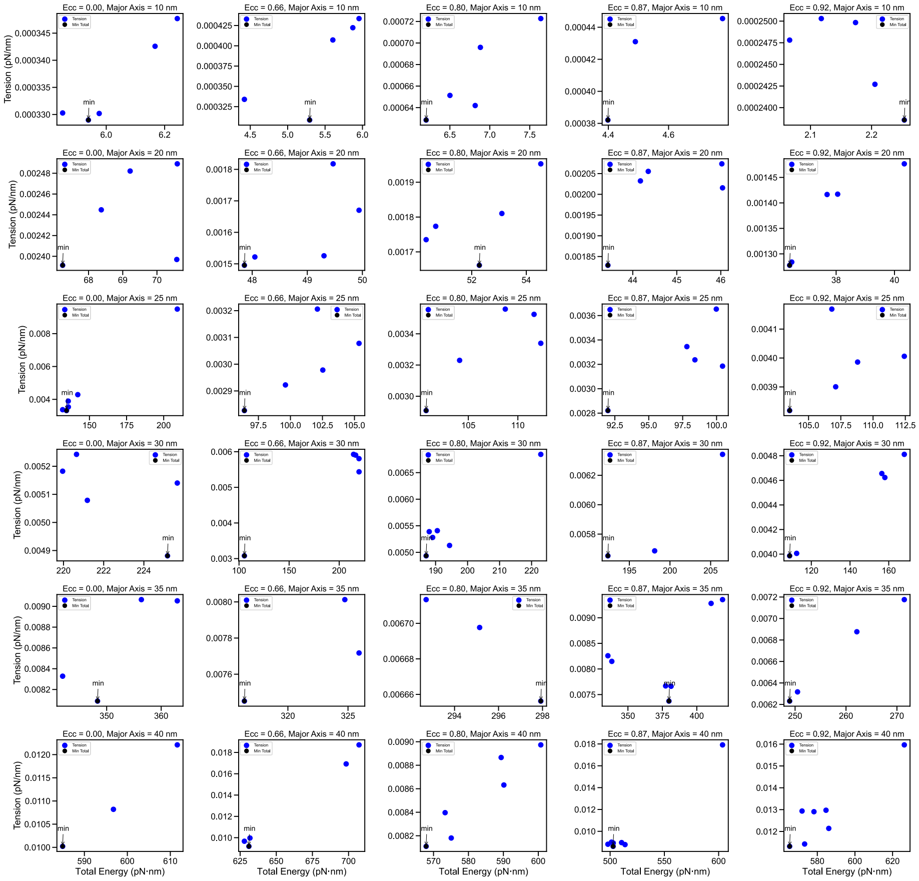

**Figure. S9.** Tension versus total energy varying eccentricity and cap size. Only samples with tension < 0.04 pN·nm are included. Each trajectory is shown as a separate data point here.

#### 2. Supporting Tables

##### 2.1. Model physical parameters

| Parameter | Definition | Value(s) in simulation |
|---|---|---|
| $\kappa$ | Membrane bending modulus | 83.4 pN·nm |
| $\mu_A$ | Area constraint constant | 250 pN/nm |
| $k_{\mathrm{hb}}$ | Harmonic bond force constant between lipid and scaffolding protein | $10^3$ pN/nm (*) |
| $C_0$ | Spontaneous curvature of membrane | 0 |
| $A_0$ | Equilibrium membrane area | 73000 $\mathrm{nm}^2$ ~ 87000 $\mathrm{nm}^2$ |
| $l_{\mathrm{b}}$ | The equilibrium harmonic bond length between lipid and scaffolding protein | 9 nm |
| $r_G$ | Spherical radius of Gag lattice sphere | 50 nm |
| $L_{\mathrm{SB}}$ | Side length of simulation box (with periodic images) | 300 nm |
| $L_{\mathrm{PB}}$ | Side length of periodic box (without periodic images) | 270 nm |
| $N_{\mathrm{sphere}}$ | Total number of Gag monomers on a fully assembled Gag lattice sphere | 3700 |
| $k_{\mathrm{B}}T$ | Boltzmann constant times temperature | 4.17 pN·nm |

**Table S1. Model physical parameters and their definitions and values used in simulation.** (*) In the simulation, $k_{\mathrm{hb}}$ is set to a small initial value $1.5\ \mathrm{pN/nm}$ and gradually increase to final value of $10^3\ \mathrm{pN/nm}$, an arbitrary large bond potential so it is effectively a rigid bond. This approach makes sure that the finite element mesh does not deform or stuck in small propagation step size due to big initial force.

#### 2.2. Model variables

| Variables | Definition | Mathematical definition |
|---|---|---|
| $E_{\text{tot}}$ | Total membrane energy | $E_{\text{tot}}(\mathbf{p}, A, r_b) \approx E_B + E_A$ |
| $E_B$ | Membrane bending energy | $E_B = \int_S \frac{1}{2}\kappa\left(2H(\mathbf{p}) - C_0(\mathbf{p})\right)^2 \sqrt{g}ds_1 ds_2$ |
| $E_A$ | Area constraint energy (global) | $E_A = \frac{1}{2}\mu_A \frac{(A - A_0)^2}{A_0}$ |
| $E_{\text{reg}}$ | Regularization energy | |
| $E_{\text{hb}}$ | Energy due to harmonic bond between lipid and scaffolding protein | $E_{\text{hb}} = \frac{1}{2}k_{\text{hb}}(r_{\text{b}} - l_{\text{b}})^2$ |
| $\mathbf{F}_{\text{hb}}$ | Force due to harmonic bond between lipid and scaffolding protein | $\mathbf{F}_{\text{hb}} = -k_{\text{hb}}(r_{\text{b}} - l_{\text{b}})\mathbf{r}_b$ |
| $E_{\text{tot,cap}}$ | Total membrane energy in the cap region | |
| $E_{\text{tot,relax}}$ | Total membrane energy in the relaxation region | |
| $n_{\text{Gag}}$ | Number of Gags in the Gag lattice | |
| $S$ | The membrane surface | |
| $\mathbf{p}$ | A point on the membrane surface | $\mathbf{p} \in S$ |
| $(s_1, s_2)$ | Curvilinear coordinates | |
| $(u, v, w)$ | Barycentric coordinates within element triangle (*) | |
| $(x, y, z)$ | Cartesian coordinates | |

| | | |
|---|---|---|
| $(\rho, \phi, h)$ | Cylindrical coordinates (azimuthal radius, azimuthal angle, height) | |
| $\mathbf{e}_1, \mathbf{e}_2$ | Tangent basis vectors | $\mathbf{e}_1 = \frac{\partial \mathbf{p}}{\partial s_1}, \mathbf{e}_2 = \frac{\partial \mathbf{p}}{\partial s_2}$ |
| $g$ | $\sqrt{g}$ is the curvilinear factor | $\sqrt{g} = \lVert \mathbf{e_1} \times \mathbf{e_2} \rVert$ |
| $H$ | Mean curvature | $H = \frac{c_1 + c_2}{2} = -\frac{(1+z_x^2)z_{yy} + (1+z_y^2)z_{xx} - 2z_x z_y z_{xy}}{2(1+z_x^2+z_y^2)^{\frac{3}{2}}}$ |
| $c_1, c_2$ | Curvature along two principal directions | |
| $A$ | Membrane surface area | |
| $\tau$ | Membrane tension | $\tau = \frac{\partial E_A}{\partial A} = \mu_A \frac{A - A_0}{A_0} = \sqrt{\frac{2\mu_A E_A}{A_0}}$ |
| $r_b$ | Distance between scaffolding protein and lipid | |
| $\mathbf{r}_b$ | Unit displacement vector from scaffolding protein to lipid | |
| $h(\rho)$ | Shape function; height in terms of azimuthal radius | |
| $r_M$ | Spherical radius of membrane cap | $r_M = r_G + l_b$ (assuming rigid and non-tilting bond) |
| $\rho_G$ | Azimuthal radius of Gag lattice cap | $\rho_G = 2r_G \sqrt{\left(\frac{n_{\text{Gag}}}{N_{\text{sphere}}}\right)\left(1 - \frac{n_{\text{Gag}}}{N_{\text{sphere}}}\right)}$ |
| $\rho_M$ | Azimuthal radius of membrane cap | $\rho_M = 2r_M \sqrt{\left(\frac{n_{\text{Gag}}}{N_{\text{sphere}}}\right)\left(1 - \frac{n_{\text{Gag}}}{N_{\text{sphere}}}\right)}$ |
| $\rho_R$ | Azimuthal radius at the end of relaxation | |

| | | |
|---|---|---|
| $\phi_R$ | Ratio of relaxation region width to membrane cap | $\phi_R = \frac{\rho_R - \rho_M}{\rho_M}$ |
| $\theta$ | Semi-vertical angle of the cap cone | $\theta = \arcsin \frac{\rho_G}{r_G}$ |
| $\mathbf{q}_l, l = 1..12$ | Coordinate vectors of 12 neighboring vertices of an element triangle (**) | |
| $\mathcal{N}_l(u, v)$ | Limit surface FEM local shape function (***) | $\mathbf{p} = \sum_{l=1}^{12} \mathcal{N}_l(u, v)\mathbf{q}_l$ |
| $\mathcal{W}(u, v)$ | Limit surface FEM global weight function | $\mathbf{p}_i = \sum_{j=1}^{m} \mathcal{W}_i(u, v)\mathbf{q}_j$ |
| $\mathbf{M}_S$ | Matrix of coordinates of membrane surface points | $\mathbf{M}_S = \begin{pmatrix} \mathbf{p}_1 \\ \mathbf{p}_2 \\ \vdots \\ \mathbf{p}_n \end{pmatrix}$ |
| $\mathbf{M}_M$ | Matrix of coordinates of control mesh vertices | $\mathbf{M}_M = \begin{pmatrix} \mathbf{q}_1 \\ \mathbf{q}_2 \\ \vdots \\ \mathbf{q}_m \end{pmatrix}$ |

**Table S2. Model physical variables and their definitions and mathematical definitions in terms of other variables or parameters.** (*) There are only two degrees of freedom in Barycentric coordinates since $u + v + w = 1$. Therefore, without loss of generality, the functions in terms of barycentric coordinates are usually written in terms of $(u, v)$. (**) Regular vertices in control mesh assumed. (***) Here "shape function" in finite element method (FEM) is different from $h(\rho)$ (function that defines membrane shape). In the field of FEM, shape functions refer to a series functions that relate the displacement at any point within the element to the position of the vertices of the element.

#### 2.3. Simulation hyperparameters

| Parameter | Definition | Value(s) used |
|---|---|---|
| `a0` | `trialStepsize = a0 * lFace * maxForceMag;` step-size resetting coefficient for Local Gradient Descent (LGD) | 0.1 |
| `lgdMagnifyCoeff` | step-size magnifying coefficient for LGD | 1.1 |
| `lgdShrinkCoeff` | step-size shrinking coefficient for LGD | 0.8 |
| `c1` | The c1 parameter in the Wolfe condition for Nonlinear Conjugate Gradient (NCG) algorithm | $10^{-6}$ |
| `c2` | The c2 parameter in the Wolfe condition | 0.001 |
| `deltaEnergyConverge` | Converging criteria for total energy | $10^{-5}$ pN·nm |
| `deltaForceScaleConverge` | Converging criteria for max force magnitude | $10^{-4}$ pN |
| `lFace` | Target side length of triangular mesh | 5 nm |
| `maxIterations` | Max number of gradient descent iterations | $10^{6}$ |
| `propagateScaffoldingInterv` | Gradient descent iteration interval for repositioning scaffolding | 7 |
| `propagateScaffoldingNstep` | Number of gradient descent steps every time propagating scaffolding | 100 |
| `updateHBConnectionNstep` | Number of gradient descent steps every time updating harmonic bond connection | 1000 |
| `stepThreshold` | Lower bound of gradient descent step size | $10^{-15}$ |

**Table S3. Hyperparameters and their values used in the simulation.** We use a nonlinear conjugate gradient (NCG) method with Wolfe Condition first and switch to simple local gradient descent (LGD) if NCG fails. See Fig. S7.

### 3. Supporting Methods

#### 3.1. Additional details on implementation of subdivision limit surface

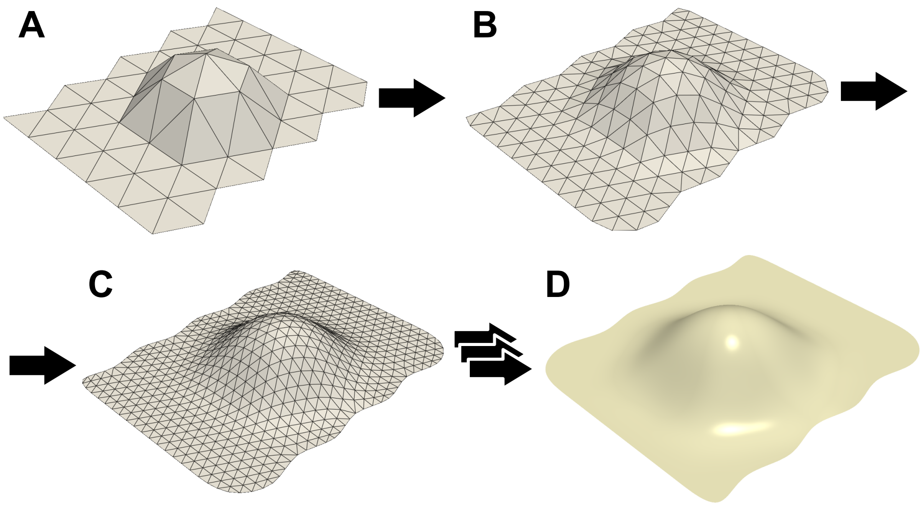

**Fig. M1. Graphic representation of the limit surface method,** which refers to gradually refining the control mesh and taking the limit to the infinite number of refinements.

##### 3.1.1. Mathematical definition of limit surface method with the shape function

As we have described in section ***II.B.1***, we numerically represent the lipid membrane surface in 3D space using a Finite Element Method (FEM) that applies the limit surface method to smoothly parameterize the surface. Here, we will elaborate on the limit surface method. Mathematically, any arbitrary point $\mathbf{p}$ on the limit surface $S$ is determined by a weighted average of the surrounding backbone vertices $\mathbf{q}_l$ of the control mesh. The point $\mathbf{p}$ can be represented by barycentric coordinates $(u, v)$ on its local element triangle, facilitating a spline fit using shape functions [1] Relative weights are given by the shape function $\mathcal{N}_l(u, v)$, where $(u, v)$ are the barycentric coordinates of point $\mathbf{p}$ on its local element triangle and $l$ denotes a neighboring backbone vertex (Fig. 1C and 1D):

$$\mathbf{p} = \sum_{l=1}^{12} \mathcal{N}_l(u,v)\mathbf{q}_l \quad (M1)$$

The full definition is the shape function $\mathcal{N}_l(u,v)$ is given by previous literature [1]. This way, we can get any point on the surface while only recording change in the mesh vertices. For a discrete approximation of the energy and force of the membrane, the model uses surface points that correspond to mesh vertices to approximate the limit surface, i.e., using characteristic points with $(u,v) = (0,0), (0,1), (1,0)$.

###### 3.1.2. Getting Conversion matrix from shape function

We define a global weight function $\mathcal{W}$ extending the domain of $\mathcal{N}_l(u,v)$ to all vertices such that it has the same value as the local shape function $\mathcal{N}_l(u,v)$ if the input vertex is one of the original 12 neighboring vertices, and outputs zero otherwise:

$$\mathcal{W}(u,v,j) = \begin{cases} \mathcal{N}_l(u,v) & (\mathbf{q}_j \in \{\mathbf{q}_l\}) \\ 0 & (\mathbf{q}_j \notin \{\mathbf{q}_l\}) \end{cases} \quad (M2)$$

This way, for some point $\mathbf{p}_i$ $(i = 1,2,\ldots,n)$, Equation M1 can be modified to iterate over all mesh points as follows. Note that $n$ is the total number of points on membrane surface and $m$ is the total number of vertices on control mesh.

$$\mathbf{p}_i = \sum_{j=1}^{m} \mathcal{W}_i(u,v)\mathbf{q}_j \quad (M3)$$

Let $\mathcal{W}_{ij} = \mathcal{W}_i(u,v,j)$. Then, as we enumerate over surface points $\mathbf{M}_S$ that correspond to each mesh vertex on the control mesh $\mathbf{M}_M$, we can calculate a conversion matrix $\mathbf{C}$ which maps between them (Equation M3). Since we have one to one correspondence between surface points and mesh points, $m = n$, $\mathbf{C}$ is a square matrix. We can increase the number of mesh point if we need higher resolution.

Commented [YY1]: Explain w_nm, compact

$$\begin{pmatrix} \mathbf{p}_1 \\ \mathbf{p}_2 \\ \vdots \\ \mathbf{p}_n \end{pmatrix} = \begin{pmatrix} \mathcal{W}_{11} & \cdots & \mathcal{W}_{1m} \\ \mathcal{W}_{21} & \cdots & \mathcal{W}_{2m} \\ \vdots & \ddots & \vdots \\ \mathcal{W}_{n1} & \cdots & \mathcal{W}_{nm} \end{pmatrix} \begin{pmatrix} \mathbf{q}_1 \\ \mathbf{q}_2 \\ \vdots \\ \mathbf{q}_m \end{pmatrix} \leftrightarrow \mathbf{M}_S = \mathbf{C}\mathbf{M}_M \tag{M4}$$

With this conversion matrix, we can calculate the matrix inverse which enables efficiently converting back and forth between the matrix of surface points $\mathbf{M}_S$ and the matrix of mesh vertices $\mathbf{M}_M$.

$$\mathbf{C}^{-1}\mathbf{M}_S = \mathbf{M}_M \tag{M5}$$

##### 3.1.3. Explicit definition of shape function in terms of barycentric coordinates

Note that because in barycentric system, $u + v + w = 1$, without loss of generality, we can write shape function in terms of $u, v$, which is $\mathcal{N}_l(u, v)$. Shape functions $\mathcal{N}_l(u, v)$ are symmetrical in terms of swapping $u, v, w$ within each of the following three groups (see Fig. M2):

1) $l = 1, 2, 6, 9, 10, 12$
2) $l = 3, 5, 11$
3) $l = 4, 7, 8$

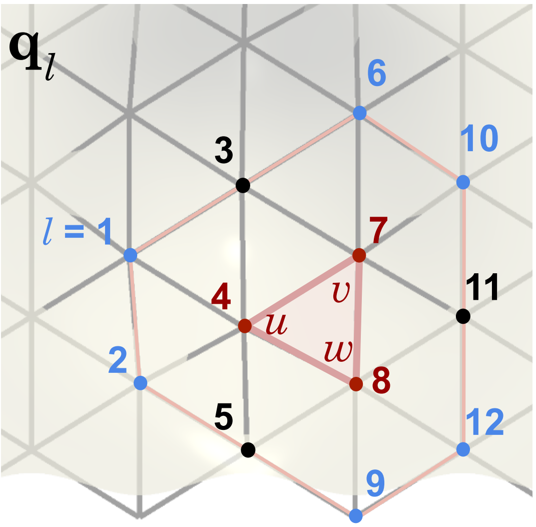

**Fig. M2.** Graphic representation of the barycentric coordinate within an element triangle and its 12 surrounding vertices.

Here, one example from each group is given as follows from previous literature [1]:

$$\begin{cases} \mathcal{N}_1(u,v) = \frac{1}{12}(u^4 + 2u^3v) \\ \mathcal{N}_3(u,v) = \frac{1}{12}(u^4 + 2u^3w + 6u^3v + 6u^2v\,w + 12u^2v^2 + 6uv^2w + 6uv^3 + 2v^3w + v^4) \\ \mathcal{N}_4(u,v) = \frac{1}{12}(6u^4 + 24u^3w + 24u^2w^2 + 8uw^3 + w^4 + 24\,u^3v + 60u^2vw + 36uvw^2 + \\ \quad 6vw^3 + 24u^2v^2 + 36uv^2w + 12v^2w^2 + 8uv^3 + 6v^3w + v^4) \\ \quad \ldots \end{cases} \tag{M6}$$

The shape function in the continuum membrane program (`src.mesh.Gauss_quadrature::get_shapefunction`) is further written as a 7 x 12 matrix, where the columns refer to the 12 neighboring vertices and the rows refer to the derivatives. This enables us to quickly get numerical values of all required derivatives of the shape function during coordinate, energy, and force calculation.

shape_functions(0,:), shape functions: $\mathcal{N}_l(u, v)$

shape_functions(1,:), differential to v: $\frac{\partial \mathcal{N}_l(u,v)}{\partial v}$

shape_functions(2,:), differential to w; $\frac{\partial \mathcal{N}_l(u,v)}{\partial w}$

shape_functions(3,:), double differential to v; $\frac{\partial^2 \mathcal{N}_l(u,v)}{\partial v^2}$

shape_functions(4,:), double differential to w; $\frac{\partial^2 \mathcal{N}_l(u,v)}{\partial w^2}$

shape_functions(5,:), differential to v and w; $\frac{\partial^2 \mathcal{N}_l(u,v)}{\partial w \partial v}$

shape_functions(6,:), differential to w and v; $\frac{\partial^2 \mathcal{N}_l(u,v)}{\partial v \partial w}$

**Table. M1.** Definition of the shape function matrix by rows. Note the first and second derivatives with respect to two of the barycentric coordinates are defined. Because the barycentric coordinates are symmetrical, we arbitrarily chose $v$ and $w$ without loss of generosity.

#### 3.2. Setup of the mesh model

##### 3.2.1. Class structure of the mesh model

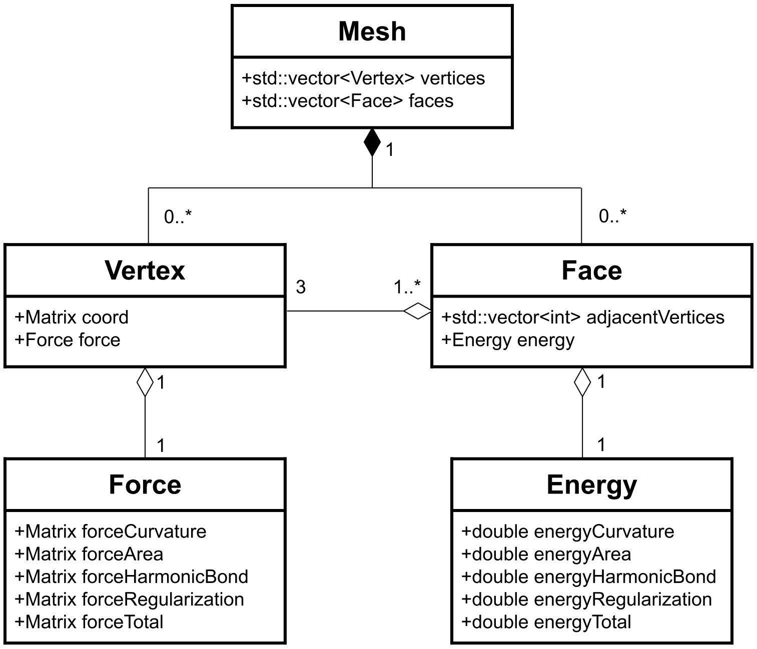

**Figure. M3. Class structure diagram of the mesh model using UML** (Unified Modeling Language). Note that the hollowed diamond arrow denotes aggregation, and the filled diamond arrow denote composition.

In the continuum membrane model, we use a simplified mesh structure without explicit definition of `edge` or `halfedge`. At the core of the class structure (Figure. M3) is the Mesh class, which aggregates a collection of `Vertex` and `Face` objects in vectors. This reflects that a mesh is composed of multiple vertices and faces, forming a structured representation of a surface. Each `Vertex` has a `coord` matrix, which stores its coordinates, and a `Force` object, responsible for modeling various forces due to membrane bending (`forceCurvature`), area stretching (`forceArea`), harmonic bond interaction between protein and membrane (`forceHarmonicBond`), and regularization (`forceRegularization`). The `Face` class represents individual element triangle and stores indices of its three adjacent vertices, along with an `Energy` object that encapsulates different energy types correspondingly.

##### 3.2.2. Geometry and Indexing setup

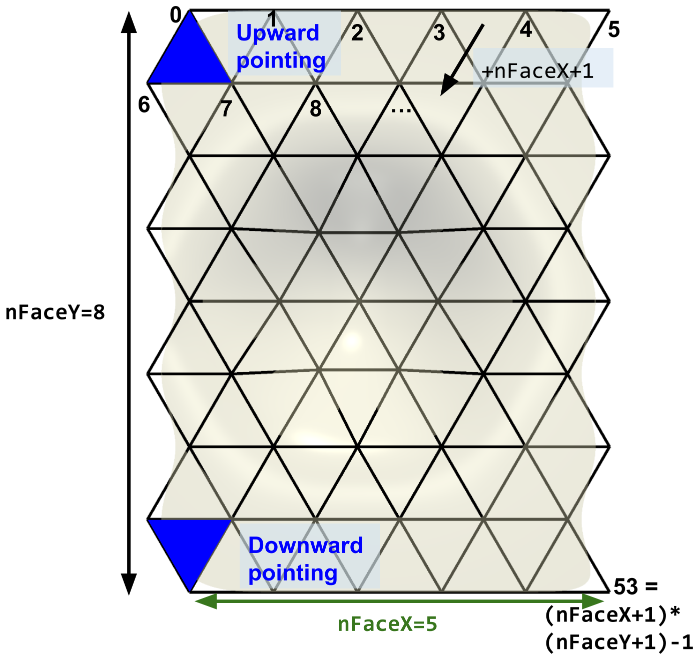

**Figure. M4. Graphic representation of the setup of a 5 x 8 mesh.** Note that the number of elements is `nFaceX * nFaceY * 2` which is 80 in the graph above and the number of vertices is `(nFaceX+1) * (nFaceY+1)` which is 54.

Here, the triangular mesh is setup with the following rules:

1. The top left triangle points upward, and the bottom left triangle points downward, which means that `nFaceY` is always even. This fixes the geometry and makes indexing easier, considering periodic boundary condition.
2. Each row has the same number of element triangles.
3. The index of vertices starts from 0 and increments first by row then by column. Row and column of vertex also starts from 0.

Now we can convert between index of vertices (`iVertex`) and (row, column) (`rVertex, cVertex`) of vertices via the following equation:

```
iVertex = rVertex * (nFaceX + 1) + cVertex
```

##### 3.2.3. Definition of face

A face is defined as an element triangle with three vertices. For any vertex (`rVertex, cVertex`) such that `rVertex ∈ [0,nFaceY]` and `cVertex ∈ [0,nFaceX]`, this vertex corresponds to two faces on the bottom right, the face index of which is assigned `(2 * iVertex)` and `(2 * iVertex + 1)` respectively, according to the figure below. Note that the sequence of vertices in the definition of faces cannot be inverted, because the sequence defines the direction of the face to distinguish between the "inside" from the "outside" of the membrane.

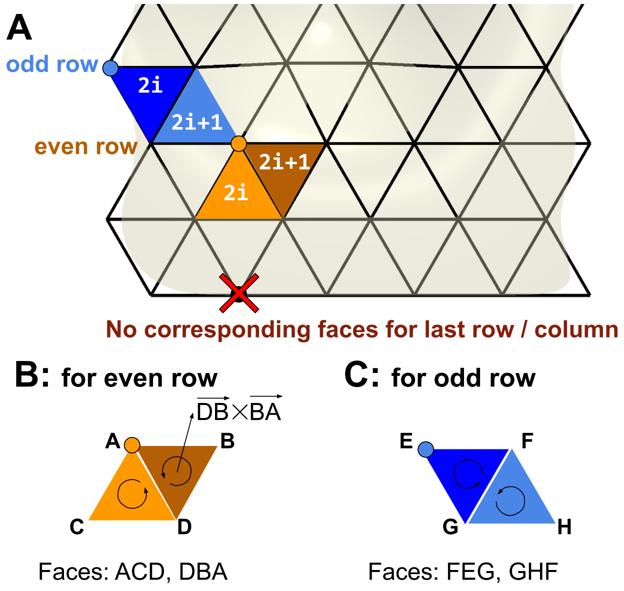

**Figure. M5.** A) Graphic representation of the two faces correspond to odd or even row vertex. B, C) The faces are defined in a specific direction such that the cross product defines the direction of face to distinguish between the "inside" from the "outside" of the membrane.

##### 3.2.4. Periodic boundary condition

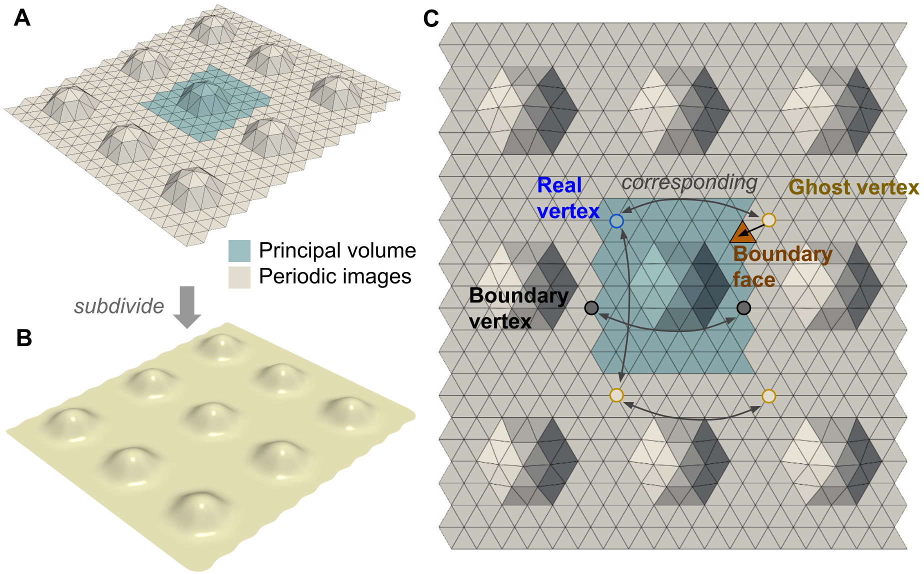

**Figure. M6. Setup of the periodic boundary condition in the simulation model.** A) The principal volume of the control mesh and its surrounding periodic images. Note that the system is only periodic in two directions (not in the direction of height). B) the subdivided limit surface with the mirror images generated from the control mesh. C) Graphical representation of real, boundary, and ghost vertices. Ghost vertices are mirror images of real vertices which help with the calculating the limit surface of boundary faces. Boundary vertices correspond to other boundary vertices and need to be updated simultaneously.

The periodic boundary is set up in two directions (excluding the direction of height) in this model (Figure. M6). The program keeps track of "ghost vertices" that are the mirror image of real vertices near the boundary to enable efficient calculation of on the boundary face, since the shape function requires the information of one ring of surrounding vertices. Note that the boundary vertices are also real but need to be treated with caution since they have pairwise correspondence and thus need to be updated simultaneously.

**Commented [YY2]:** Call it periodic images instead of mirror images since they are not mirror symmetry

##### 3.3. Analytical solution to relaxation shape function that minimizes bending energy

The problem analytical solving the shape function of the detachment region of the membrane attached to a sphere has been investigated by various previous research [2]. However, a few assumptions must be made to enable an analytical or numerical solution. Here, the following cases are examined:

| **Assumption** | Ignorable tension | Small gradient approximation |
|---|---|---|
| **Section 3.3.1.** | Yes | Yes |
| **Section 3.3.2.** | Yes | No |
| **Section 3.3.3.** | No | Yes |

**Statement of the problem**: Let $(\rho, \phi, h)$ be cylindrical coordinates and $(x, y, z)$ be the corresponding cartesian coordinates and assume the system is in azimuthal symmetry. Now consider bending energy $E_B$:

$$E_B = \int_x \int_y 2\kappa \left(H(x,y)\right)^2 dydx \tag{M7}$$

With the boundary conditions with respect to the shape function:

- Continuity at transition between cap and relaxation: $h(\rho_M) = 0$
- Differentiability at transition between cap and relaxation: $h'(\rho_M) = -\tan\theta$
- Membrane becomes flat at the end of relaxation: $h'(\rho_R) = 0$

What is the shape function $h(\rho)$ such that the membrane bending energy is minimized, when the membrane is attached to a perfect spherical cap with distance $r_b$ and goes to flat at infinite distance assuming azimuthal symmetry?

##### 3.3.1A. Derivation of analytical solution with small gradient approximation

Here, $(x, y, z)$ are Eucleadian coordinates and $(\rho, \phi, h)$ are cylindrical coordinates. In our model with small gradient approximation, we have $z_x^2$ << 1, $z_y^2$ << 1:

$$H\big(\mathbf{p}(x, y, z)\big) = -\frac{(1 + z_x^2)z_{yy} + \big(1 + z_y^2\big)z_{xx} - 2z_x z_y z_{xy}}{2\big(1 + z_x^2 + z_y^2\big)^{\frac{3}{2}}} \approx -\frac{1}{2}(\nabla^2 z) \qquad (\text{M8})$$

Plug in zero spontaneous curvature ($C_0(\mathbf{p}) = 0$) and express bending energy in terms of x, y, z:

By using cylindrical coordinates defined by:

$$\begin{cases} x = \rho\cos\phi \\ y = \rho\sin\phi \\ z = h \end{cases}$$

Therefore, equation M7 can be further expressed in cylindrical coordinates as:

$$E_B = \int_\rho \int_{\phi=0}^{2\pi} \frac{1}{2}\kappa\left(\frac{1}{\rho}\frac{\partial}{\partial\rho}\left(\rho\frac{\partial h}{\partial\rho}\right)\right)^2 \rho d\phi d\rho = \pi\kappa\int_\rho \frac{1}{\rho}\left(\frac{\partial}{\partial\rho}\left(\rho\frac{\partial h}{\partial\rho}\right)\right)^2 d\rho$$

Because of radio symmetry, the height is only dependent on radius. Therefore, the equation above can be expressed in total derivative:

$$E_B = \pi\kappa\int\frac{1}{\rho}\left(\frac{d}{d\rho}\left(\rho\frac{dh}{d\rho}\right)\right)^2 d\rho = \pi\kappa\int\frac{1}{\rho}(h' + \rho h'')^2 d\rho \qquad (\text{M9})$$

The minimum of Equation M9 can be solved using Euler-Lagrange method with the integrand defined as:

$$L(\rho, h, h', h'') = \rho\left(\frac{h'}{\rho} + h''\right)^2 \qquad \text{(M10)}$$

Plug in Euler-Lagrange Equation $\frac{\partial L}{\partial h} - \frac{d}{d\rho}\left(\frac{\partial L}{\partial h'}\right) + \frac{d^2}{d\rho^2}\left(\frac{\partial L}{\partial h''}\right) = 0$, the simplified Euler-Lagrange equation is then:

$$\frac{d}{d\rho}\left(\frac{h'}{\rho} + h''\right) = \frac{d^2}{d\rho^2}(h' + \rho h'') \qquad \text{(M11)}$$

Simplify and we get a Euler-Cauchy equation that can be easily solved:

$$\rho^3 h'''' + 2\rho^2 h''' - \rho h'' + h' = 0 \qquad \text{(M12)}$$

To correctly represent the logarithmic operations in the solution to the Euler-Cauchy equation, we first put our system in dimensionless form by setting the characteristic length to $\rho_M$. Namely, for some length $l$, the dimensionless length $\bar{l}$ is defined by the following equation:

$$\bar{l} = \frac{l}{\rho_M} \qquad \text{(M13)}$$

The Euler-Cauchy equation now becomes:

$$\bar{\rho}^3 \frac{d^4\bar{h}}{d\bar{\rho}^4} + 2\bar{\rho}^2 \frac{d^3\bar{h}}{d\bar{\rho}^3} - \bar{\rho}\frac{d^2\bar{h}}{d\bar{\rho}^2} + \frac{d\bar{h}}{d\bar{\rho}} = 0$$

Solving the equation, we get (where $c_i$ are real constants):

$$\tilde{h}(\rho) = c_1\rho^2 + c_2\rho^2 \ln(\rho) + c_3 \ln(\rho) + c_4 \tag{M14}$$

Note that we have more constants than boundary conditions, which means that we have an infinite number of solutions. Therefore, we need to search in the parameter space such that the bending energy is minimized. Note the derivatives:

$$\begin{cases} \tilde{h} = c_1\rho^2 + c_2\rho^2 \ln\rho + c_3 \ln\rho + c_4 \\ \dfrac{d\tilde{h}}{d\rho} = 2c_1\rho + c_2(\rho + 2\rho\ln\rho) + \dfrac{c_3}{\rho} \\ \dfrac{d^2\tilde{h}}{d\rho^2} = 2c_1 + c_2(3 + 2\ln\rho) - \dfrac{c_3}{\rho^2} \end{cases} \tag{M15}$$

Recall bending energy with small gradient approximation. Note that $\frac{E_B}{\pi\kappa}$ is dimensionless,

$$\frac{E_B}{\pi\kappa} = \int_{\rho_M}^{\rho_R} \frac{1}{\rho}(h' + \rho h'')^2 d\rho = \int_1^{\tilde{\rho}_R} \frac{1}{\rho}\left(\frac{d\tilde{h}}{d\rho} + \rho\frac{d^2\tilde{h}}{d\rho^2}\right)^2 d\rho$$

Replace the derivatives and we have,

$$\frac{E_B}{\pi\kappa} = \int_1^{\tilde{\rho}_R} \frac{1}{\rho}\left(2c_1\rho + c_2(\rho + 2\rho\ln\rho) + \frac{c_3}{\rho} + \rho\left(2c_1 + c_2(3 + 2\ln\rho) - \frac{c_3}{\rho^2}\right)\right)^2 d\rho$$

$$= 16\int_1^{\tilde{\rho}_R} \rho(c_1 + c_2 + c_2\ln\rho)^2 d\rho$$

Then, dimensionless bending energy can be simplified to:

$$\frac{E_B}{\pi\kappa} = 16\int_1^{\tilde{\rho}_R} \rho(c_1 + c_2 + c_2\ln\rho)^2 d\rho = 16(c_1 + c_2)^2 \int_1^{\tilde{\rho}_R} \rho\left(1 + \frac{c_2}{c_1 + c_2}\ln\rho\right)^2 d\rho$$

Integrate and expand the expression, we have:

$$\frac{E_B}{\pi\kappa} = 4(c_1 + c_2)^2 \left( (\tilde{\rho_R}^2 - 1) \left( 2 - 2\left(\frac{c_2}{c_1 + c_2}\right) + \left(\frac{c_2}{c_1 + c_2}\right)^2 \right) \right.$$

$$\left. - 2\left(\frac{c_2}{c_1 + c_2}\right)\left(\frac{c_2}{c_1 + c_2} - 2\right)\tilde{\rho_R}^2 \ln\tilde{\rho_R} + 2\left(\frac{c_2}{c_1 + c_2}\right)^2 \tilde{\rho_R}^2 (\ln\tilde{\rho_R})^2 \right)$$

Simplify,

$$\frac{E_B}{\pi\kappa} = 2\left((\tilde{\rho_R}^2 - 1)((2c_1 + c_2)^2 + c_2^2) + 4c_2(2c_1 + c_2)\tilde{\rho_R}^2 \ln\tilde{\rho_R} + 4c_2^2 \tilde{\rho_R}^2 (\ln\tilde{\rho_R})^2\right) \quad (\text{M16})$$

At this point, reconsider the three boundary conditions. Boundary condition 1 (membrane is continuous at transition) basically translates the membrane up and down and is thus related to $c_4$. Therefore, focus on Boundary condition 2 and 3 and plug in the expression for $h'$, we have:

$$\begin{cases} -\tan\theta = 2c_1 + c_2 + c_3 \\ 0 = 2c_1\tilde{\rho_R} + c_2(\tilde{\rho_R} + 2\tilde{\rho_R}\ln\tilde{\rho_R}) + \dfrac{c_3}{\tilde{\rho_R}} \end{cases}$$

Eliminate $c_3$ by substitution, we get:

$$-\tan\theta = 2c_1 + c_2 - 2c_1\tilde{\rho_R}^2 - c_2(\tilde{\rho_R}^2 + 2\tilde{\rho_R}^2 \ln\tilde{\rho_R})$$

Simplify, we get:

$$2c_1 + c_2 = \frac{\tan\theta - 2c_2(\tilde{\rho_R}^2 \ln\tilde{\rho_R})}{\tilde{\rho_R}^2 - 1} \quad (\text{M17})$$

Here, we simplify equation M17 above such that $2c_1 + c_2 = u + vc_2$ by grouping up the constants with $u, v \in \mathbb{R}$ defined by:

$$\begin{cases} u = \dfrac{\tan\theta}{\widetilde{\rho_R}^2 - 1} \\ v = -\dfrac{2\left(\widetilde{\rho_R}^2 \ln\widetilde{\rho_R}\right)}{\widetilde{\rho_R}^2 - 1} \end{cases} \tag{M18}$$

Note that because $\rho_R > \rho_M$, $\widetilde{\rho_R}^2 - 1 > 0$. And thus $u > 0$ and $v < 0$. Plug $2c_1 + c_2 = u + vc_2$ back into equation M16, we get:

$$\frac{E_B}{\pi\kappa} = 2\left(\left(\widetilde{\rho_R}^2 - 1\right)\left((u + vc_2)^2 + c_2^2\right) + 4c_2(u + vc_2)\widetilde{\rho_R}^2 \ln\widetilde{\rho_R} + 4c_2^2 \widetilde{\rho_R}^2 \left(\ln\widetilde{\rho_R}\right)^2\right) \tag{M19}$$

Note that the dimensionless bending energy is a quadratic with respect to $c_2$, i.e. for fixed relaxation region size $\widetilde{\rho_R}$, there exist real constants $\alpha_0, \alpha_1, \alpha_2$ such that the dimensionless bending energy can be expressed as:

$$\frac{E_B}{\pi\kappa} = \alpha_2 c_2^2 + \alpha_1 c_2 + \alpha_0$$

Note that the quadratic term scaler $\alpha_2$ is positive as:

$$\begin{aligned} \alpha_2 &= 2\left(\widetilde{\rho_R}^2 - 1\right)\left(v^2 + 1\right) + 8v\widetilde{\rho_R}^2 \ln\widetilde{\rho_R} + 8\widetilde{\rho_R}^2 \left(\ln\widetilde{\rho_R}\right)^2 \\ &= 2\left(\widetilde{\rho_R}^2 - 1\right)\left(v^2 + 1\right) + 8\widetilde{\rho_R}^2 \ln\widetilde{\rho_R}\left(v + \ln\widetilde{\rho_R}\right) \end{aligned}$$

Note that $v + \ln\widetilde{\rho_R} = \frac{\left(\widetilde{\rho_R}^2 - 1\right)\ln\widetilde{\rho_R} - 2\left(\widetilde{\rho_R}^2 \ln\widetilde{\rho_R}\right)}{\widetilde{\rho_R}^2 - 1} - \ln\widetilde{\rho_R}\frac{\widetilde{\rho_R}^2 + 1}{\widetilde{\rho_R}^2 - 1}$

$$\begin{aligned} \alpha_2 &= 2\left(\widetilde{\rho_R}^2 - 1\right)\left(\left(\frac{2\left(\widetilde{\rho_R}^2 \ln\widetilde{\rho_R}\right)}{\widetilde{\rho_R}^2 - 1}\right)^2 + 1\right) - 8\widetilde{\rho_R}^2 \left(\ln\widetilde{\rho_R}\right)^2 \frac{\widetilde{\rho_R}^2 + 1}{\widetilde{\rho_R}^2 - 1} \\ &= -\frac{8\widetilde{\rho_R}^2 \left(\ln\widetilde{\rho_R}\right)^2}{\widetilde{\rho_R}^2 - 1} + 2\left(\widetilde{\rho_R}^2 - 1\right) \end{aligned}$$

**Statement:**

$f(x) > 0$ for all $x > 1$, where real function $f(x) = -4x^2(\ln x)^2 + (x^2-1)^2$.

**Proof:**

Use monotonicity theorem. Let real function $g(y) = e^{2y} - 2y^2 - 2y - 1$ defined on $y \in \mathbb{R}$. At $y = 0$, $\frac{dg}{dy} = 2e^{2y} - 4y - 2 = 2 - 0 - 2 = 0$. Again, $\forall y > 0$,

$$\frac{d^2g}{dy^2} = 4e^{2y} - 4 > 0$$

Therefore, $\forall y > 0, \frac{dg}{dy} > 0$. Similarly, because $g(0) = 0$, $\forall y > 0, g > 0$. Let $x = e^y$, then

$$y > 0 \Leftrightarrow x > 1$$

Therefore, the statement $\forall y > 0, g > 0$ is equivalent to:

$$\forall x > 1, x^2 - 2(\ln x)^2 - 2\ln x - 1 > 0$$

Now consider the derivative $\frac{df}{dx}$. $\forall x > 1$,

$$\frac{df}{dx} = 4x(x^2 - 2(\ln x)^2 - 2\ln x - 1) > 0$$

Since at $x = 1$, $f(1) = 0 + 0 = 0$, again with monotonicity theorem, $\forall x > 1$

$$f(x) = -4x^2(\ln x)^2 + (x^2-1)^2 > 0$$

By statement above, because $\widetilde{\rho_R}^2 > 1$,

$$\alpha_2 = \frac{2}{\widetilde{\rho_R}^2 - 1}\left(-4\widetilde{\rho_R}^2 \ln \widetilde{\rho_R}^2 + \left(\widetilde{\rho_R}^2 - 1\right)^2\right) > 0$$

And therefore, this quadratic function reaches its minimum at $c_2 = -\frac{\alpha_1}{2\alpha_2}$. Simplify and we get:

$$c_2 = -\frac{4\left(\rho_{\overline{R}}^{\ 2} - 1\right)uv + 4u\rho_{\overline{R}}^{\ 2}\ln\rho_{\overline{R}}}{4\left(\rho_{\overline{R}}^{\ 2} - 1\right)\left(v^2 + 1\right) + 16v\rho_{\overline{R}}^{\ 2}\ln\rho_{\overline{R}} + 16\rho_{\overline{R}}^{\ 2}\left(\ln\rho_{\overline{R}}\right)^2}$$

Substitute the definition $v = -\frac{2\left(\rho_{\overline{R}}^{\ 2}\ln\rho_{\overline{R}}\right)}{\rho_{\overline{R}}^{\ 2}-1}$ into the nominator of the equation above, we see:

$$c_2 = -\frac{-4u\left(\rho_{\overline{R}}^{\ 2} - 1\right)\frac{2\left(\rho_{\overline{R}}^{\ 2}\ln\rho_{\overline{R}}\right)}{\rho_{\overline{R}}^{\ 2} - 1} + 4u\rho_{\overline{R}}^{\ 2}\ln\rho_{\overline{R}}}{4\left(\rho_{\overline{R}}^{\ 2} - 1\right)\left(v^2 + 1\right) + 16v\rho_{\overline{R}}^{\ 2}\ln\rho_{\overline{R}} + 16\rho_{\overline{R}}^{\ 2}\left(\ln\rho_{\overline{R}}\right)^2} = 0$$

Therefore, we conclude that to minimize $E_B$, $c_2$ must be equal to zero. Hence, the shape function in this case becomes $\hbar(\rho) = c_1\rho^2 + c_3\ln(\rho) + c_4$. The three boundary conditions can be expressed as:

$$\begin{cases} \hbar(1) = c_1 + c_4 = 0 \\ \hbar'(1) = 2c_1 + c_3 = -\tan\theta \\ \hbar'(\rho_{\overline{R}}) = 2c_1\rho_{\overline{R}} + \frac{c_3}{\rho_{\overline{R}}} = 0 \end{cases}$$

Solving the system of equations above, we get the constants of the dimensionless shape function:

$$\begin{cases} c_1 = \frac{\tan\theta}{\rho_{\overline{R}}^{\ 2} - 1} \\ c_3 = -2\rho_{\overline{R}}^{\ 2}\frac{\tan\theta}{\rho_{\overline{R}}^{\ 2} - 1} \\ c_4 = -\frac{\tan\theta}{\rho_{\overline{R}}^{\ 2} - 1} \end{cases} \tag{M20}$$

Lastly, we introduce back the units and recover the shape function,

$$\frac{h(\rho)}{\rho_M} = \frac{c_1\rho^2}{\rho_M^{\ 2}} + c_3\ln\left(\frac{\rho}{\rho_M}\right) + c_4 \Longrightarrow h(\rho) = \frac{c_1\rho^2}{\rho_M} + c_3\rho_M\ln\left(\frac{\rho}{\rho_M}\right) + c_4\rho_M \tag{M21}$$

The dimensionless bending energy corresponding to the shape function above is then,

$$\frac{E_B}{\pi\kappa} = 8\left(\left(\tilde{\rho_R}^2 - 1\right)c_1^2\right) = \frac{8\tan^2\theta}{\tilde{\rho_R}^2 - 1} \tag{M22}$$

And thus, the corresponding bending energy is:

$$E_B = \frac{8\pi\kappa\tan^2\theta}{\tilde{\rho_R}^2 - 1} = 8\pi\kappa\tan^2\theta\,\frac{\rho_M^2}{\rho_R^2 - \rho_M^2} \tag{M23}$$

Here we see the observation that bending energy goes to zero as relaxation radius goes to infinity,

$$\lim_{\tilde{\rho_R}\to\infty} E_B = 0$$

##### 3.3.1B. Analogy of Michell Solution to the analytical solution with small gradient approximation

The Michell Solution [3] refers to a general solution to the elasticity equation as stated below in terms of Airy stress function $\psi$.

$$\nabla^4\psi = 0$$

With the following boundary requirement: let $\frac{\partial\psi}{\partial x} = H(x, y) + \alpha$ and $\frac{\partial\psi}{\partial y} = K(x, y) + \beta$ where $H, K$ are real functions of $x, y$ and $\alpha, \beta$ are real constants, we have:

(1) $\psi$ of any point on the surface is equal to the values that integrates from the boundary following $H$ and $K$.

(2) $\psi$ and its derivatives go back to their starting value when integrated along the path of the boundary

Due to the similarity of the problem stated in section S4.1, where the equation for minimum bending energy conformation is in the following form with the same requirement on the boundary, we can apply the same solution in this scenario (with an extra constant term $q_0$ to shift the height based on the placement of cylindrical coordinate system).

$$\nabla^4 h = 0$$

Let $(\rho, \theta, h)$ be the cylindrical coordinates. The Michell solution is in the form of a Fourier series in $\theta$:

$$h(\rho, \theta) = a_0\rho^2 + b_0\rho^2 \ln\rho + c_0 \ln\rho + q_0 + \theta \cdot I(\rho) + \sum_{n=1}^{\infty}\left(\cos(n\theta) \cdot G(\rho, n) + \sin(n\theta) \cdot H(\rho, n)\right)$$

Where $a_0, b_0, c_0, q_0$ are real constants and $G, H, I$ are real functions. Note that, in the scenario defined in section S4.1, we assume azimuthal symmetry around $\theta$. In this case the solution simply becomes:

$$h(\rho) = a_0\rho^2 + b_0\rho^2 \ln\rho + c_0 \ln\rho + q_0 \quad \text{(M24)}$$

Which conforms to our solution in section 3.3.1A.

###### 3.3.2. Numerical shape function solution without small gradient approximation

With greater deviation from plane, the small gradient approximation no longer. In this case, starting again from the following equation, let us consider the same problem still with zero spontaneous curvature but without small gradient approximation (SGA). The curvature is now expressed as,

$$H(\mathbf{p}(x, y, z)) = -\frac{(1 + z_x^2)z_{yy} + (1 + z_y^2)z_{xx} - 2z_x z_y z_{xy}}{2(1 + z_x^2 + z_y^2)^{\frac{3}{2}}} \quad \text{(M25)}$$

To express the equation above in cylindrical coordinate system (note the height remains unchanged $z = h$), let's consider the derivatives:

$$\begin{cases} z_x = h_\rho \cos\phi - \frac{h_\phi \sin\phi}{\rho} \\ z_y = h_\rho \sin\phi + \frac{h_\phi \cos\phi}{\rho} \end{cases} \tag{M26}$$

Assuming radial symmetry, now we can write height $h$ purely in terms of azimuthal radius $\rho$. Note $h^{(n)} = \frac{d^n h}{(d\rho)^n}$ for some positive integer $n$:

$$\begin{cases} z_x = h' \cos\phi \\ z_y = h' \sin\phi \\ z_{xx} = h'' \cos^2\phi + \frac{h' \tan^2\phi}{\rho(1+\tan^2\phi)} \\ z_{yy} = h'' \sin^2\phi + \frac{h'}{\rho(1+\tan^2\phi)} \\ z_{xy} = z_{yx} = h'' \sin\phi \cos\phi - \frac{h' \tan\phi}{\rho(1+\tan^2\phi)} \end{cases} \tag{M27}$$

Substitute equation M27 to equation M25 and now we can simplify the curvature equation to:

$$H(\rho) = -\frac{h'' + \frac{h'}{\rho} + \frac{h'^3}{\rho}}{2(1+h'^2)^{\frac{3}{2}}} \tag{M28}$$

Using the result above, the bending energy equation M7 then turns into:

$$E_B = \int_\rho \int_{\phi=0}^{2\pi} \frac{1}{2}\kappa \frac{\left(h'' + \frac{h'}{\rho} + \frac{h'^3}{\rho}\right)^2}{(1+h'^2)^3} \rho d\phi d\rho = \pi\kappa \int \frac{\left(h'' + \frac{h'}{\rho} + \frac{h'^3}{\rho}\right)^2}{(1+h'^2)^3} \rho d\rho \tag{M29}$$

We have Euler-Lagrange Equation $\frac{\partial L}{\partial h} - \frac{d}{d\rho}\left(\frac{\partial L}{\partial h'}\right) + \frac{d^2}{d\rho^2}\left(\frac{\partial L}{\partial h''}\right) = 0$, where $L(\rho, h, h', h'') = \frac{\rho\left(h'' + \frac{h'}{\rho} + \frac{h'^3}{\rho}\right)^2}{(1+h'^2)^3}$. We use the following Mathematica code (`solve_EL_full_wo_SGA.nb`) to simplify the Euler-Lagrange Equation:

```
(*Define the function f*)
f[x_, a_, b_, c_] := x*(c + b/x + b^3/x)^2/(1 + b^2)^3

(*Replace variables*)
fReplaced = f[\[Rho], h[\[Rho]], h'[\[Rho]], h''[\[Rho]]]

(*Compute partial derivatives*)
dfdb = D[fReplaced, h'[\[Rho]]]  (*Partial derivative w.r.t b=h'[rho]*)
dfdc = D[fReplaced,
  h''[\[Rho]]] (*Partial derivative w.r.t c=h''[rho]*)

(*Compute the required derivatives*)
ddfdbdrho = D[dfdb, \[Rho]]    (*First derivative of dfdb w.r.t rho*)
d2dfdcdrho2 =
 D[dfdc, {\[Rho], 2}] (*Second derivative of dfdc w.r.t rho*)

(*Display results*)
eqn = Simplify[ddfdbdrho - d2dfdcdrho2]
```

The output equation from Mathematica to solve is:

$$
\begin{aligned}
0 = -\frac{1}{\rho^2(1+h'[\rho]^2)^5} 2 \Big( & 3h'[\rho]^5 + h'[\rho]^7 + 3\rho h'[\rho]^6 h''[\rho] \\
& + h'[\rho]\big(1 - 9\rho^2 h''[\rho]^2 - 12\rho^3 h''[\rho] h^{(3)}[\rho]\big) \\
& - 3h'[\rho]^3\big(-1 + 3\rho^2 h''[\rho]^2 + 4\rho^3 h''[\rho] h^{(3)}[\rho]\big) \\
& + \rho h'[\rho]^4 \Big(5h''[\rho] + \rho\big(2h^{(3)}[\rho] + \rho h^{(4)}[\rho]\big)\Big) \\
& + \rho \Big(-h''[\rho] - 3\rho^2 h''[\rho]^3 + \rho\big(2h^{(3)}[\rho] + \rho h^{(4)}[\rho]\big)\Big) \\
& + \rho h'[\rho]^2 \Big(h''[\rho] + 21\rho^2 h''[\rho]^3 + 2\rho\big(2h^{(3)}[\rho] + \rho h^{(4)}[\rho]\big)\Big)\Big)
\end{aligned}
$$

To further solve the equation, we rewrite the equation in terms of $h^{(4)}$ (fourth derivative of $h$) on the left-hand-side:

$$h^{(4)}[\rho] = -\frac{1}{\rho^3(1+h'[\rho]^2)^2}(3h'[\rho]^5 + h'[\rho]^7 + 3\rho h'[\rho]^6 h''[\rho] - \rho(h''[\rho] + 3\rho^2 h''[\rho]^3 - 2\rho h^{(3)}[\rho]) + \rho h'[\rho]^4(5h''[\rho] + 2\rho h^{(3)}[\rho]) + \rho h'[\rho]^2(h''[\rho] + 21\rho^2 h''[\rho]^3 + 4\rho h^{(3)}[\rho]) + h'[\rho](1 - 9\rho^2 h''[\rho]^2 - 12\rho^3 h''[\rho]h^{(3)}[\rho]) - 3h'[\rho]^3(-1 + 3\rho^2 h''[\rho]^2 + 4\rho^3 h''[\rho]h^{(3)}[\rho]))$$

Now, we can use numerical solver to solve the equation above. Note that in addition to the three boundary conditions, we require a fourth boundary condition because this is now a fourth order ODE.

$$\begin{cases} h(\rho_M) = 0 \\ h'(\rho_M) = -\tan\theta \\ h'(\rho_R) = 0 \\ h''(\rho_M) = h''_{\text{init}} \end{cases}$$

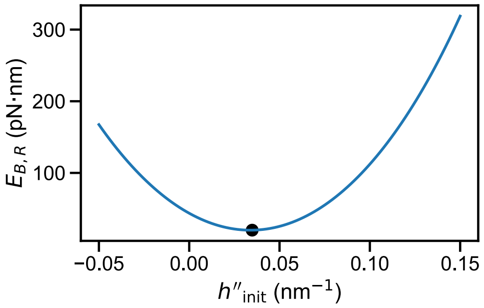

**Figure. M7. A range of different initial second derivatives of height and their corresponding bending energy of relaxation region.** Through this method, we can get the initial second derivative such that the bending energy is minimized (shown in back dot on the figure).

Because we are looking for the solution such that the bending energy is minimized, we iterate through a range of $h''_{\text{init}}$ and plot $h''_{\text{init}}$ against bending energy to search for the minimum with the following python code (see Fig. M7). Here, we use boundary value problem (BVP) solvers and numerical integrators in `scipy.integrate` module.

```
import numpy as np
from scipy.integrate import solve_bvp

def ode_system(x, y):
    """Defines the system of differential equations to solve."""
    h, h1, h2, h3 = y  # Unpacking state variables
    rho = x  # Radial coordinate

    # Define h'''' from the governing equation
    f = (-1 / (rho**3 * (1 + h1**2)**2)) * (3 * h1**5 + h1**7 + 3 * rho * h1**6 * h2
           - rho * (h2 + 3 * rho**2 * h2**3 - 2 * rho * h3) + rho * h1**4 * (5 * h2 + 2 * rho * h3)
           + rho * h1**2 * (h2 + 21 * rho**2 * h2**3 + 4 * rho * h3) + h1 * (1 - 9 * rho**2 * h2**2 - 12 * rho**3 * h2 * h3)
           - 3 * h1**3 * (-1 + 3 * rho**2 * h2**2 + 4 * rho**3 * h2 * h3))

    # Return system of ODEs
    return [y[1], y[2], y[3], f]

def bc(ya, yb, init_slope = -0.1, init_secondderiv = 0.04):
    """Defines the boundary conditions for the system."""
    return np.array([ya[0], ya[1] - init_slope, ya[2] - init_secondderiv, yb[1]])

# Initialize solution array
rhoM = 20.0
rhoR = 40.0
# Define the radial range for solving the BVP
x = np.linspace(rhoM, rhoR, 1000)
y_a = np.zeros((4, x.size))
y_a[1] = 0.0  # Initial guess for the first derivative

# Solve the boundary value problem
res_a = solve_bvp(ode_system, bc, x, y_a)
```

We define a dimensionless ratio $\phi_R$ to characterize the ratio between azimuthal radius of relaxation and azimuthal radius of membrane cap:

$$\phi_R = \frac{\rho_R - \rho_M}{\rho_M} \tag{M30}$$

Like the simulations we did in the main paper, here we calculate the relationship between minimum bending energy versus the size of the scaffolding lattice while fixing $\phi_R$ (the graph below shows the result for $\phi_R = 1$). We see that relaxation almost grows linearly with respect to the number of scaffolding nodes.

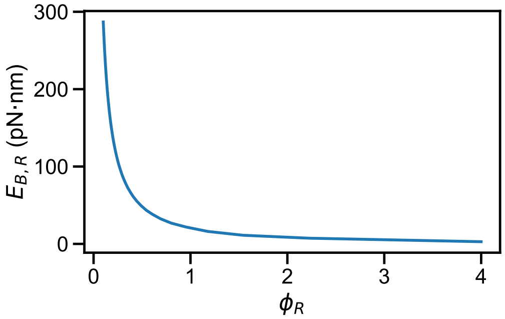

**Figure. M8. The bending energy of relaxation area consistently goes down to approach zero as we increase the azimuthal radius of relaxation region without small gradient approximation.** The analytical solution in plot uses the following parameters: $\rho_M = 20$ nm, $r_M = 59$ nm.

If we repeat this process now with fixed cap geometry (number of scaffolding nodes and eccentricity), we can plot the relationship between bending energy of the relaxation area ($E_{B,R}$) and $\phi_R$ shown in the following figure. As we see here, the bending energy of relaxation area still consistently goes down to approach zero as we increase the azimuthal radius of relaxation region, even in the scenario without small gradient approximation.

##### 3.3.3. Analytical shape function considering tension with small gradient approximation

One major shortcoming of the shape function solution above is that it completely ignores the contribution of tension. Here, we use constrained Euler-Lagrange equation to solve for best configuration while considering nonzero tension. Since we have total energy:

$$E = E_B + E_A \tag{M31}$$

Let $h^*(\rho)$ be some shape function and $E_B^*$ be the bending energy of the minimum bending energy state constraining the total membrane area to $A^*$, then because $E_A$ is only related to total area, $h^*$ is also the shape function such that the total energy $E^*$ is minimized constraining the total membrane area to $A^*$. Therefore, for the set of all possible total membrane area $\{A^*\}$, we have a corresponding set of total energy $\{E^*\}$ and the overall minimum total energy is reached at $\min\{E^*\}$ (Here assuming $A = A_0$). Therefore, the global area constraint can be approximately described by:

$$A^* = 2\pi \int_{\rho_M}^{\rho_R} \rho\sqrt{1 + h'^2}\, d\rho \approx \pi(\rho_R^2 - \rho_M^2) + \pi \int_{\rho_M}^{\rho_R} \rho h'^2\, d\rho \tag{M32}$$

Let $\hbar(\rho) = h'(\rho)$ then we have constraint, $\int \rho\hbar^2 d\rho = C$ where $C$ is a constant defined by $C = \frac{A^*}{\pi} - (\rho_R^2 - \rho_M^2)$. Thus, let $\mathcal{F}(\rho) = \rho\hbar^2(\rho) - \frac{C}{\rho_M - \rho_R}$, we have:

$$\int_{\rho_M}^{\rho_R} \mathcal{F}(\rho) d\rho = \int_{\rho_M}^{\rho_R} \rho\hbar^2(\rho) d\rho - C = 0 \tag{M33}$$

Use constrained Euler-Lagrange equation $\frac{d}{d\rho}\left(\frac{\partial L}{\partial \hbar'}\right) - \frac{\partial L}{\partial \hbar} = -\lambda \frac{\partial \mathcal{F}}{\partial \hbar}$ , where we defined the Euler-Lagrange function as follows:

$$L(\rho, \hbar, \hbar') = \rho \left( \frac{\hbar}{\rho} + \hbar' \right)^2 \quad \text{(M34)}$$

Plug in and simplify the equation, we get:

$$\rho^2 \hbar'' + \rho \hbar' - \hbar + \lambda \rho^2 \hbar = 0 \quad \text{(M35)}$$

Which can be solved analytically to get:

$$\hbar(\rho) = c_5' J_1(\sqrt{\lambda}\rho) + c_6' Y_1(\sqrt{\lambda}\rho) \quad \text{(M36)}$$

where $J_{\mathrm{n}}$ is Bessel function of the first kind and $Y_n$ is the Bessel function of the second kind; $c_5', c_6'$ are real constants. Since $\hbar = h'$, the shape function of relaxation region is then:

$$h(\rho) = c_5 J_0(\sqrt{\lambda}\rho) + c_6 Y_0(\sqrt{\lambda}\rho) + c_7 \quad \text{(M37)}$$

where $\sqrt{\lambda}, c_5, c_6, c_7$ are parameters. By using the same 3 boundary conditions in section II.A.4 and the global area constraint defined in Equation above, we can solve for $\sqrt{\lambda}, c_5, c_6, c_7$. However, solving the following system of equations requires a numerical solver. By numerically solving the equation fixing the total membrane area at a range of different values, we can plot total energy against surface area of the membrane. Taking the minimum point on the total energy curve yields the minimum total energy state. This result further reiterates the point that directly setting $E_A = 0$ is completely different from allowing lipid to freely diffuse in and out of system boundary i.e. allowing $A_0$ to freely change without any energy penalties, even though the theoretical minimum area constraint energy at the end is both zero. In the limit of an infinite boundary, this result is identical to the previously published result. Again, by numerically solving the equation fixing the total membrane area at a range of different values, we can plot bending, area stretching, and total energy against surface area of the membrane. The result is shown below.

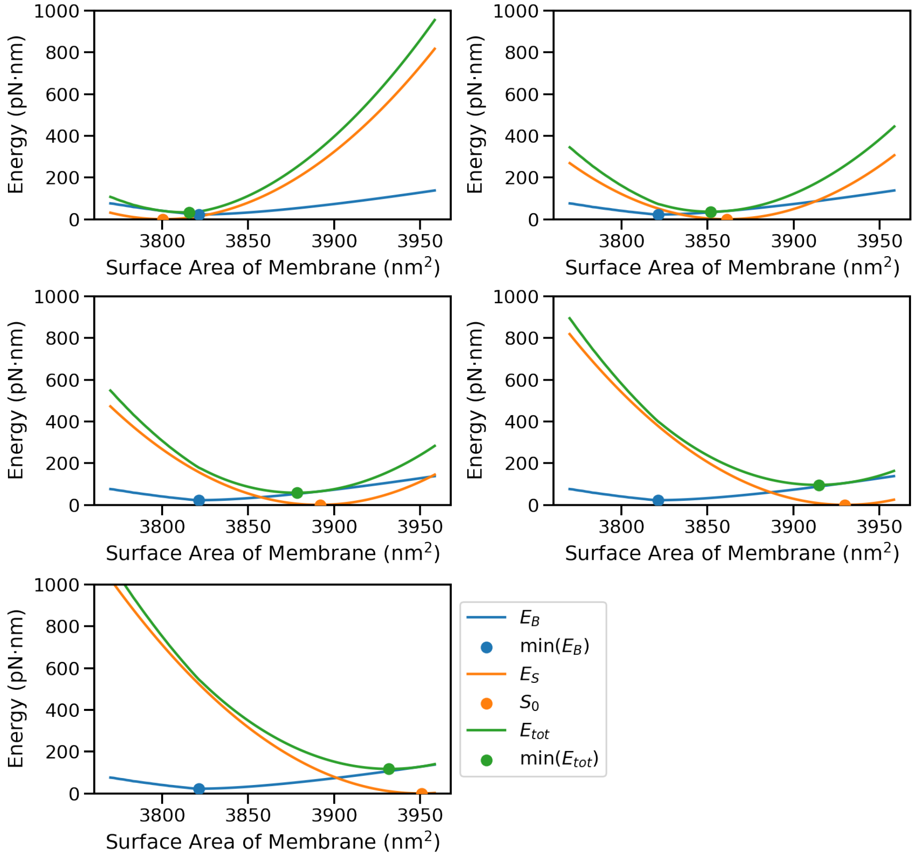

**Figure. M9. The minimum bending energy, area constraint energy, and total energy of the relaxation region when varying the surface area of the membrane. Multiple cases are shown**

**changing the relax area $S_0$ and thus shifting the area constraint energy curve.** The analytical solution in plot uses the following parameters: $\rho_M = 20\ \mathrm{nm}, \rho_R = 40\ \mathrm{nm}, r_M = 59\ \mathrm{nm}$.

We see that here, when shifting the area constraint energy, the surface area at which total energy reaches minimum always lies closer to the minimum point of area constraint energy curve. However, at total energy minimum, area constraint energy always has a much smaller contribution to the total energy than bending energy. This can be explained mathematically. Suppose we have some energy function $\mathcal{E}$ which consists of two quadratic functions $\mathcal{E}_{1,2}$ with respect to the same variable $x$. Both quadratic functions have the property that the function equals to zero at minimum, i.e.

$$\mathcal{E}(x) = \mathcal{E}_1(x) + \mathcal{E}_2(x) = \mu_1(x - x_1)^2 + \mu_2(x - x_2)^2 \tag{M38}$$

Where $\mu_{1,2}$ are positive real constants and $\mu_1 > \mu_2 > 0$. $x_{1,2}$ are the $x$ at which the corresponding function yields minimum. Now consider the minimum of $\mathcal{E}(x)$.

$$\mathcal{E}(x) = (\mu_1 + \mu_2)x^2 - 2(\mu_1 x_1 + \mu_2 x_2)x + \mu_1 x_1^2 + \mu_2 x_2^2 \tag{M39}$$

$\mathcal{E}(x)$ yields minimum at $x = x_{\mathcal{E}}$:

$$x_{\mathcal{E}} = \frac{\mu_1 x_1 + \mu_2 x_2}{\mu_1 + \mu_2} \tag{M40}$$

Which is a convex combination between $x_1$ and $x_2$, and thus $x_{\mathcal{E}}$ is closer to $x_1$ as $\mu_1 > \mu_2 > 0$. This also means that changing $x_1$ causes a bigger shift in $x_{\mathcal{E}}$ compared to changing $x_2$. Now we plug in to get the corresponding energies:

$$\begin{cases} \mathcal{E}_1(x_\mathcal{E}) = \mu_1 \left( \frac{\mu_1 x_1 + \mu_2 x_2}{\mu_1 + \mu_2} - x_1 \right)^2 = \mu_1 \left( \frac{\mu_2}{\mu_1 + \mu_2} (x_2 - x_1) \right)^2 \\ \mathcal{E}_2(x_\mathcal{E}) = \mu_2 \left( \frac{\mu_1 x_1 + \mu_2 x_2}{\mu_1 + \mu_2} - x_2 \right)^2 = \mu_2 \left( \frac{\mu_1}{\mu_1 + \mu_2} (x_1 - x_2) \right)^2 \end{cases} \quad \text{(M41)}$$

It is obvious that $\mathcal{E}_1(x_\mathcal{E}) < \mathcal{E}_2(x_\mathcal{E})$ since

$$\frac{\mathcal{E}_1(x_\mathcal{E})}{\mathcal{E}_2(x_\mathcal{E})} = \frac{\mu_1 \mu_2^2}{\mu_1^2 \mu_2} < 1 \quad \text{(M42)}$$

Therefore, $\mathcal{E}_1(x_\mathcal{E})$ contributes less to the energy, even though $x_\mathcal{E}$ is closer to $x_1$. This explains the phenomenon we see in Fig. M9 that the surface area at which total energy reaches minimum always lies closer to the minimum point of area constraint energy curve, but at total energy minimum, area constraint energy always has a much smaller contribution to the total energy than bending energy.

##### 3.3.4. Circumference approximated by surface area versus by binding sites

To establish a relationship between the azimuthal radius of a spherical membrane cap $\rho_M$ and the number of Gag monomers on the cap $n_{\mathrm{Gag}}$, a naïve way is to assume that each Gag monomer takes up the same surface area on the membrane cap and thus we can calculate the membrane cap surface area $A_{\mathrm{cap}}$ from $n_{\mathrm{Gag}}$:

$$A_{\mathrm{cap}} = 2\pi r_M h_{\mathrm{cap}} = n_{\mathrm{Gag}} \bar{A}_{\mathrm{Gag}} \tag{M43}$$

Where $h_{\mathrm{cap}}$ is the height of the spherical cap and $N_{\mathrm{Gag}}$ is the theoretical number of Gag monomers to fully assemble a Gag lattice sphere (here we set to $N_{\mathrm{Gag}} = 3700$ for $r_M = 59\ \mathrm{nm}$). From this expression, calculating $\rho_M$ becomes trivial:

$$\rho_M = f_1(n_{\mathrm{Gag}}) = \sqrt{2r_M h_{\mathrm{cap}} - h_{\mathrm{cap}}^2} = \frac{n_{\mathrm{Gag}} \bar{A}_{\mathrm{Gag}}}{2\pi r_M} \sqrt{\frac{4\pi r_M^2}{n_{\mathrm{Gag}} \bar{A}_{\mathrm{Gag}}} - 1} \tag{M44}$$

The problem with this model $\rho_M = f_1(n_{\mathrm{Gag}})$ is that it overestimates the size of spherical cap because there is a discrepancy between circumference calculated by surface area and circumference surrounding all the binding sites. The distance between those two circumferences $l_{\mathrm{hg}}$ is approximately half of the Gag distance $l_g$:

$$l_{hg} = \frac{1}{2} l_g$$

From that, we can construct a geometric model and calculate the circumference surrounding all the binding sites according to Figure 4.3 below.

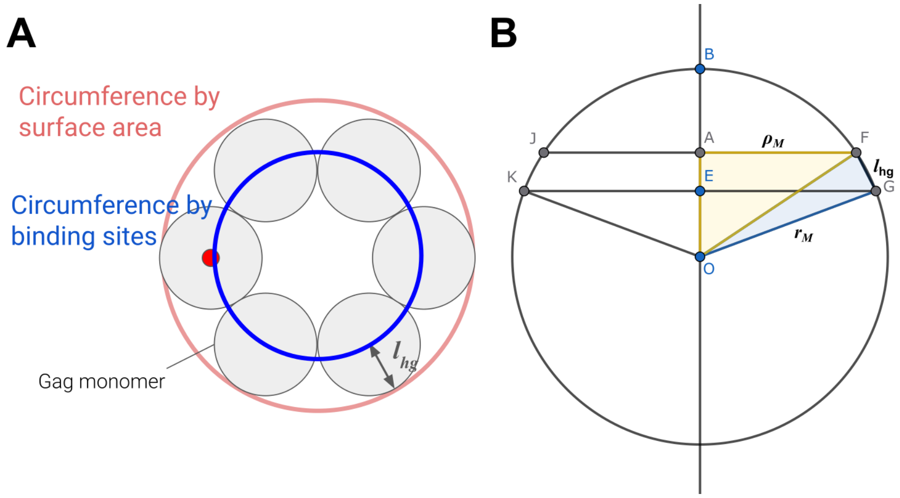

**Fig. M10.** A) Discrepancy between circumference calculated by surface area and circumference surrounding all the binding sites and the distance is approximately half of the distance between Gags; B) geometric model to calculate the circumference surrounding all the binding sites.

Given some number of Gags $n_{\text{Gag}}$, suppose the circumference defined by surface area stops at $G$ whereas the circumference defined by the outskirt of binding sites stops at $F$. Here, we need to solve for $\rho_M = \|AF\|$. Consider triangle $EOG$ and $FOG$, we have:

$$\begin{cases} \cos\angle EOG = 1 - \dfrac{\|BE\|}{r_M} = 1 - \dfrac{n_{\text{Gag}} \bar{A}_{\text{Gag}}}{2\pi r_M^2} \\ \cos\angle FOG = \dfrac{2r_M^2 - l_{hg}^2}{2r_M^2} = 1 - \dfrac{l_{hg}^2}{2r_M^2} \end{cases}$$

Therefore,

$$\rho_M = f_2(n_{\text{Gag}}) = \|OF\| \sin\angle AOF = r_M(\sin\angle EOG \cdot \cos\angle FOG - \cos\angle EOG \cdot \sin\angle FOG)$$

The difference between model $f_1$ and $f_2$ is shown in the following figure.

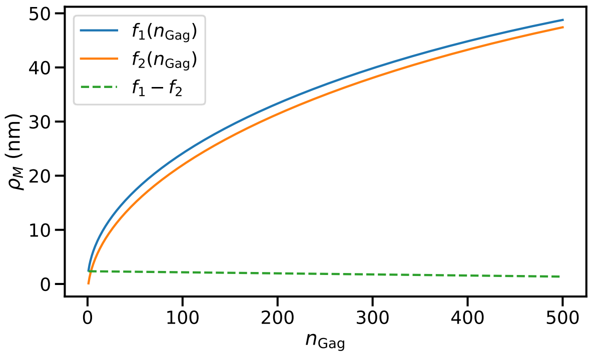

**Fig. M11. The relationship between number of Gags and azimuthal radius of Gag lattice cap model with $f_1$ versus $f_2$.** The difference between the two models slightly decays as number of Gags increases.

From the plot, we see that there is a very slow and almost linear decay of difference in azimuthal radius $\Delta\rho_M = f_1(n_{\mathrm{Gag}}) - f_2(n_{\mathrm{Gag}})$ from $f_1(0) - f_2(0)$. Here we quantify this difference.

```
Circumference_approximated_by_surface_area_vs_binding_sites.ipynb
```

##### 3.4. Theoretical lattices from Monte Carlo optimization

To establish a theoretical baseline to compare with the self-assembled lattices from NERDSS simulation, a Monte Carlo optimization program is used to generate theoretical Gag lattices with given lattice size and eccentricity. Here, the Gag monomers are simplified to nodes lying on the spherical surface, fully ignoring the orientation of interaction. Specifically, the Monte Carlo optimization starts with a set of randomly placed nodes representing the Gag monomers on the Gag spherical surface within boundary.

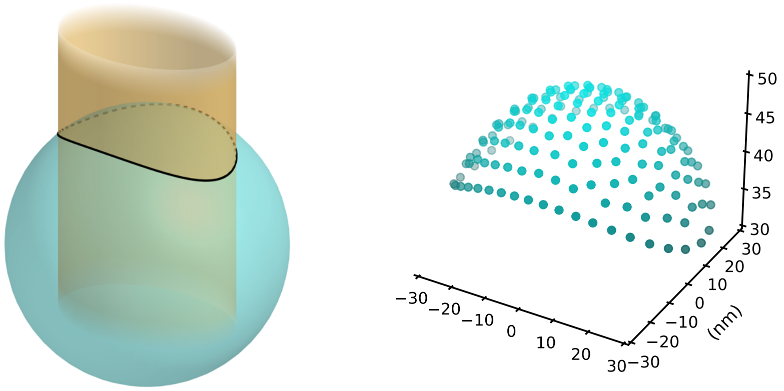

**Fig. M12. Ideal irregular vertices are set up by getting the area on a spherical shell enclosed with an elliptical pipe.** We can then use the eccentricity of the cross-section ellipse as a representation of irregularity.

Even if the Gag monomers are simplified to nodes lying on the spherical surface, fully ignoring the orientation of interaction, generating an "ideal" mesh that has "evenly placed nodes" that are $l_{\mathrm{gg}} = 4.7\ \mathrm{nm}$ (average Gag-Gag interaction length) [4] apart within a particular geometric boundary can be challenging, due to the topological layout of the nodes rely strongly on the shape of the surface. To generate an ideal mesh that has evenly placed nodes, we used a Monte-Carlo simulation to start with a given number of nodes, and let the nodes propagate on the surface based on Lennard-Jones Potential, with the minimum Lennard-Jones energy at where nodes are exactly $l_{\mathrm{gg}}$ apart. To accelerate the simulation, we apply simply the summed Lennard-Jones potential of the system to only calculate the potential between each node and their closest neighbor using KDTree.

However, there is one known drawback of this algorithm: this method currently may not correctly place nodes on the boundary of highly eccentric lattices, which is shown in section 3.4.4.

##### 3.4.1. Estimating the number of nodes for a given cap size and shape

This algorithm, however, requires us to provide the number of nodes as input parameter. To achieve that, we provide the program with a starting number of nodes and let it relax. If the

This problem is thus stated as: given the bounded surface defined by the equation below,

$$\begin{cases} x^2 + y^2 + z^2 = r^2 \\ \dfrac{x^2}{a^2} + \dfrac{y^2}{b^2} \leq 1 \end{cases}$$

where without loss of generality, $0 < a \leq b \leq r$, estimate the maximum number of nodes that can be evenly placed with each node being $l_{gg}$ apart from its closest neighbor on this surface.

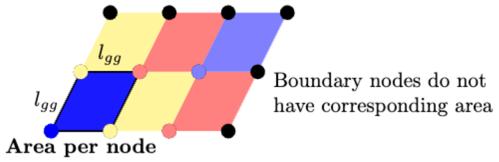

**Fig. M13.** Only half of the boundary nodes edges have corresponding area.

We first use equilateral triangle mesh to approximate the placement of nodes in the densest configuration as required by the statement. The area per node is the area of the parallelogram above. With Gag-Gag interaction length estimated as 4.7 nm [4], the area per node is:

$$\text{Area per node} = \frac{\sqrt{3}}{2} l_{\text{gg}}^2 \approx 19.13\,\text{nm}^2$$

The number of nodes is approximated by the following equation, where the first term corresponds to the nodes with corresponding parallelograms in the figure above and the second term correspond to the nodes without corresponding parallelograms. Thus, the number of nodes can be estimated as:

$$\text{Number of nodes} \approx \frac{\text{Area}_{\text{up}}}{\text{Area per node}} + \frac{N_{\text{boundary}}}{2}$$

Note that because we are on a spherical rather than perfectly flat surface, plus the fact that the is a harsh boundary condition, the surface cannot be tiled with perfect equilateral triangles. Thus, the number of nodes above is usually an overestimate of the number of nodes.  To illustrate, from the figure above, approximately half of the boundary points do not have corresponding parallelogram area. Using the approximation $\text{Circumference}_{\text{up}} \approx \pi(a+b)$

, we get:

$$N_{\text{boundary}} \leq \left\lfloor \frac{\text{Area}_{\text{up}}}{\text{Area per node}} \right\rfloor + \frac{\text{Circumference}_{\text{up}}}{2l_{gg}} \approx \left\lfloor \frac{\text{Area}_{\text{up}}}{\text{Area per node}} \right\rfloor + \left\lfloor \frac{\pi(a+b)}{2l_{gg}} \right\rfloor$$

Convert the equations that defines the boundary to spherical coordinates:

$$\begin{cases} x = r\sin\theta\cos\phi \\ y = r\sin\theta\sin\phi \\ z = r\cos\theta \end{cases}$$

where $0 \leq \theta \leq \frac{\pi}{2}, 0 \leq \phi \leq 2$ and r is a constant because we are on a spherical surface.

From the definition of the boundary surface, the equation for the boundary is:

$$\frac{x^2}{r^2} + \frac{y^2}{r^2} + \frac{z^2}{r^2} = 1 = \frac{x^2}{a^2} + \frac{y^2}{b^2}$$

This simplifies to:

$$z^2 = \frac{x^2}{a^2} + \frac{y^2}{b^2} - \frac{x^2}{r^2} - \frac{y^2}{r^2}$$

Define $c^2 = \frac{r^2}{a^2} - 1, \quad d^2 = \frac{r^2}{b^2} - 1$, where $c > 0, d > 0$. Because $a, b, r$ are constants, $c, d$ are also constants. By plugging c, d into the equation above and writing the equation in spherical coordinate, we have:

$$\cos^2\theta = c^2\sin^2\theta\cos^2\phi + d^2\sin^2\theta\sin^2\phi$$

Note that $cot\,\theta \geq 0$ as $0 \leq \theta \leq \frac{\pi}{2}$, we get the equation for the boundary:

$$\cot\theta = \sqrt{c^2\cos^2\phi + d^2\sin^2\phi}$$

The area of the upper surface $\text{Area}_{\text{up}}$ is given by:

$$\text{Area}_{\text{up}} = r^2 \int_0^{2\pi} \int_0^{\theta_0(\phi)} \sin\theta\, d\theta d\phi = r^2 \int_0^{2\pi} \left(1 - \cos\theta_0(\phi)\right) d\phi$$

Note that for any real number α, $\cos(\cot^{-1}(\alpha)) = \frac{1}{\sqrt{\frac{1}{\alpha^2}+1}}$. Therefore, the area of the upper surface $\text{Area}_{\text{up}}$ can be simplified to:

$$\text{Area}_{\text{up}} = r^2 \int_0^{2\pi} \left( 1 - \frac{1}{\sqrt{\frac{1}{c^2 \cos^2 \phi + d^2 \sin^2 \phi} + 1}} \right) d\phi \qquad (\text{M48})$$

In the special case where $c = d = 0$, the bounded surface becomes a hemisphere cap. The upper surface area becomes $\text{Area}_{\text{up}} = 2\pi r^2$, conforming to the geometry. By using the following code, we get the Gag lattice mesh with ideal perimeter shapes.

##### 3.4.2. Pseudo-code and example trajectory of the algorithm

```
1. Define class `ParticlePlacementSimulation` to perform particle placement
simulation.

 1.1. Define `__init__`:
   - Initialize parameters for simulation:
   - Sphere radius, ellipse dimensions, desired particle density, Lennard-Jones
potential parameters and node management variables.
   - Use the upper surface area calculated from the equation above to compute the
initial number of particles (`N`).

 1.2. Define `approximate_spherical_cap_area`:
   - Compute the surface area of an elliptical cap on a sphere using the equation for
Area_up above with numerical intergration.

 1.3. Define `calculate_lj`:
   - Calculate the Lennard-Jones potentials between two particles.
   - Calculate the Lennard-Jones force vector between two particles.

 1.4. Define `constrain`:
    - Constrain particle positions to the simulation's boundary conditions.

 1.5. Define `monte_carlo_random_move`:
    - Perform Monte Carlo random displacement of particles.

 1.6. Define `check_convergence`:
    - Check if the simulation has converged based on Lennard-Jones potential history.

2. Define `run_simulation`:
```

- Initialize particle positions randomly on the sphere.
- Iteratively compute forces, move particles, and adjust parameters.
- Use adaptive step sizes and Monte Carlo random moves for optimization.
- Check for convergence and adjust the number of particles if necessary.

Example trajectory of generating an ideal mesh:

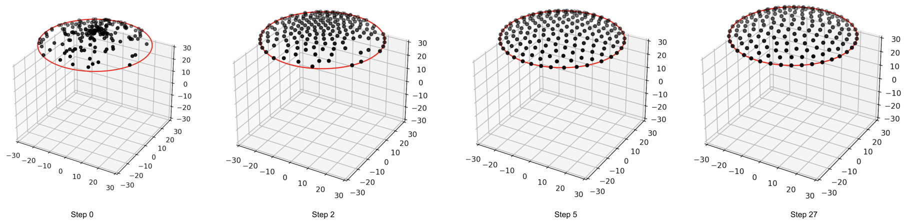

**Fig. M14. Example trajectory of generating Gag lattice mesh with ideal perimeter shapes implementing the pseudo-code above.** The points representing center of mass protein monomers are randomly placed on within the cap at the start (step 0) and pushed apart via Lennard-Jones Potential.

##### 3.4.3. Using eccentricity to evaluate irregularity of ideal Gag lattice

As mentioned above, we use a lattice in the shape of an ellipse cutting through a spherical cap. Therefore, we can use the eccentricity of the ellipse to establish a theoretical scale for evaluating irregularity of ideal Gag lattice. For an ellipse defined as:

$$\frac{x^2}{a^2} + \frac{y^2}{b^2} = 1, (a \geq b) \quad (M49)$$

Where $a$ is the major axis and $b$ is the minor axis. The eccentricity $e$ is then defined as:

$$e(a, b) = \sqrt{1 - \frac{b^2}{a^2}} \quad (M50)$$

Note that the eccentricity of a circle equals 0 and the eccentricity of a parabola equals 1. Therefore, as the eccentricity of the lattices goes up, the lattice becomes more irregular, and the range of our interest is $e \in [0,1)$.

##### 3.5. Bootstrap statistics of minimum

In this research, we use bootstrap statistics to calculate the confidence interval of minimum and to systematically determine whether there are statistically significant differences in the minimum in the energies. The scripts for bootstrap statistics in the following section are uploaded to GitHub repository.

###### 3.5.1. Resampling test of minimum

The resampling test of minimum estimates the error of the minimum within the dataset. First, within each Gag distance group, the available energy values are randomly resampled with replacement to create new synthetic datasets, where each resampled dataset has the same number of values as the original group. This process is repeated 100,000. For each resampled dataset, the minimum energy value is recorded and appended to the dataset of resampled minimum, within which a one-sided confidence bound at the specified confidence level (value used in this work: 95%) is calculated.

###### 3.5.2. Joint resampling test with respect to minimum total energy

The joint resampling test is a bootstrapping approach based on resampling test designed to preserve the relationships between physical variables in a dataset. In each bootstrap iteration, entire rows of the dataset are randomly selected with replacement, keeping all the values in a row linked as they were in the original data. From each resampled dataset, the row with the smallest total energy is chosen, and all the other associated values from that row (such as bending energy and tension) are recorded.

Repeating this process many times builds up distributions of these values under resampling. A one-sided confidence interval is then computed for total energy, while two-sided intervals are calculated for the other quantities (since other quantities are not at their corresponding minima, rather we are sampling those quantities such that the total energy is minimized).

##### 3.5.3. Permutation test of difference between minima

The permutation test starts with calculating the observed difference in the minimum values of the two datasets. Then, the two datasets are pooled into a single array and shuffled. In each iteration, the shuffled data is split back into two groups of the same original sizes, and the difference in the minimum values of the two resampled groups is recorded. The p-value is then calculated as the proportion of times this resampled difference is equal to or greater than the observed difference.

#### 3.6. Open-source code for simulation and graphs

The source code related to this work is fully open sourced on GitHub under GNU public license v3.0.

##### 3.6.1. Open-source continuum membrane model

https://github.com/mjohn218/continuum_membrane

(Placeholder, ^ this link is the private development repo for continuum membrane)

##### 3.6.2. Open-source code for the graphs and analytical and numerical calculations

https://github.com/yingyue2030699/continuum-membrane-si

(Placeholder, ^ this link is also currently the private development repo. We will either publish it or make a new public one)

##### 3.6.3. Open-source code for NERDSS (Nonequilibrium Reaction-diffusion Self-assembly Simulator)

Placeholder.com

### 4. References

#### 4.1. Softwares and packages referenced in this work

| |
|---|
| C++ Version 14<br>• OpenMP 6.0<br>• GNU Scientific Library 2.8<br>• GoogleTest 1.16.0 |
| Python 3.9.7<br>• Numpy 1.26.4<br>• Pandas 1.3.4<br>• Scipy 1.13.1<br>• PyVista 0.38.3<br>• Matplotlib 3.7.0<br>• Seaborn 0.12.1<br>• Celluloid 0.2.0<br>• Graphviz 0.20.3 |
| Mathematica 14.0.0.0 (Mac OS X ARM (64-bit)) |
